# Hippocampal theta frequency as a readout of path-integration recalibration

**DOI:** 10.64898/2026.05.06.723266

**Authors:** Seong-Beom Park, Manu S. Madhav, Ravikrishnan P. Jayakumar, Noah J. Cowan, James J. Knierim

**Affiliations:** Zanvyl Krieger Mind/Brain Institute, Johns Hopkins University, Baltimore, MD, 21218, USA; Laboratory for Computational Sensing and Robotics, Johns Hopkins University, Baltimore, MD, 21218, USA; Kavli Neuroscience Discovery Institute, Johns Hopkins University, Baltimore, MD, 21218, USA; Department of Mechanical Engineering, Johns Hopkins University, Baltimore, MD, 21218, USA; Solomon H. Snyder Department of Neuroscience, Johns Hopkins University, Baltimore, MD, 21205, USA

## Abstract

Understanding how the brain represents hidden variables is a fundamental challenge. In navigation, the internal path-integration gain is often masked by external landmarks that override the path integrator. Path integration can recalibrate its gain when allothetic and idiothetic cues conflict, but the real-time dynamics of this process are hidden to direct observation. Here, we demonstrate that theta frequency provides an error signal between the observable hippocampal gain and the internal path-integration gain. Theta frequency decreased as conflict between landmark-driven hippocampal gain and path-integration gain increased and recovered as the path-integration gain recalibrated to the new gain. A continuous attractor model replicated these dynamics, suggesting that the theta-frequency drop is driven by the misalignment of allothetic and idiothetic inputs, reducing the excitatory drive to the network. Thus, theta frequency provides a real-time readout of internal gain-error signals, offering a novel methodology to estimate hidden cognitive variables through observable physiological oscillations.

## Introduction

One of the central challenges in cognitive neuroscience is to understand how the brain processes internal representations of variables that cannot be directly manipulated or observed^1, 2^ (e.g., internal representations of confidence, reward prediction, or path-integration-based position^3–5^). Because these internal variables are often inaccessible to direct observation, researchers must rely on indirect behavioral and physiological readouts to infer their presence and dynamics. This challenge becomes especially difficult when these internal variables exhibit continuous change over time.

Spatial navigation provides a prime opportunity to explore this question. According to cognitive map theory, the hippocampus constructs an internal representation of the environment that supports flexible navigation and memory-guided behavior in animals and episodic memory in humans^6^. Both allothetic sensory cues (such as visual landmarks) and path-integration computations (which rely on idiothetic signals, such as vestibular and proprioceptive input, to estimate position relative to previous locations) work together to build and update this map^6, 7^. In familiar, stable environments, allothetic landmarks prevent or correct the cumulative errors that result from imprecisions in the path integrator system. Accordingly, allothetic landmarks will usually override the ongoing path integration system, making it difficult to separate the contributions of allothetic and idiothetic signals under naturalistic conditions. Traditional approaches to studying path integration perform experiments in the dark to minimize the influence of allothetic landmarks^8–13^. As a consequence, these approaches are limited in their ability to test direct interactions between path integration and allothetic landmark navigation during most navigation tasks, when both sets of cues contribute substantially and simultaneously^14–16^.

To overcome this limitation, we developed a virtual reality environment in which animals can move through physical space while visual landmarks are dynamically adjusted in real time^17–19^. Jayakumar and colleagues^17^ recorded CA1 place fields in this environment as rats ran on a circular track while visual landmarks moved either in the same direction as the rat or in the opposite direction with a speed proportional to the rat’s movement speed. In most sessions, the *hippocampal gain*—the factor relating movement through space to the updating of position on the hippocampal cognitive map—was tightly locked to the visual gain. After many laps, the visual landmarks were removed, allowing the path-integration dynamics to be investigated in the absence of the overriding influence of the salient landmarks. The results showed that the *path-integration gain*—i.e. the factor relating self-motion cues to the rate of movement through the hippocampal map—had been partially recalibrated during the preceding laps during which landmarks were in conflict with self-motion cues. However, because of the dominance of the landmarks, the dynamics of path-integration gain recalibration could not be directly measured during the recalibration process itself when the landmarks were present. This raises the question: is it possible to infer the hidden path-integration gain from hippocampal activity recorded while landmarks were still visible?

In this study, we present a novel method to infer the path-integration gain by utilizing two components: the directly observable hippocampal gain and an error signal that quantifies the current mismatch between the hippocampal gain and the path-integration gain. One promising candidate for such an error signal is the hippocampal theta rhythm, a prominent 7-9 Hz oscillation observed in the hippocampal local field potential (LFP)^20–23^. Theta frequency may reflect the mismatch between external sensory input and internal expectations^24, 25^, but it is unknown whether it quantitatively codes the *magnitude* of prediction error and whether it can represent the mismatch between hippocampal gain and path-integration gain while landmarks are present. To test this, we analyzed LFP data^17^ to examine whether recalibration of path integration is correlated with changes in hippocampal theta frequency.

## Results

### Theta frequency decreases during experimental gain change but increases when the gain change stops

In the previous study^17^, rats moved around the perimeter of a circular table while tethered to a radial arm that permitted only counterclockwise movement along a circular trajectory (Fig. 1a). Three visual landmarks, projected onto the interior surface of a surrounding hemispherical dome structure, were rotated around the center of the environment as a function of the rat’s movement; this rotation produced a continuous mismatch between idiothetic estimates of position and visual landmark feedback. The direction and magnitude of landmark rotation were determined by an experimental gain, *G*, defined as the distance traveled by the rat in the virtual reference frame defined by the landmarks divided by the distance traveled in the physical reference frame of the laboratory.

**Figure 1.**
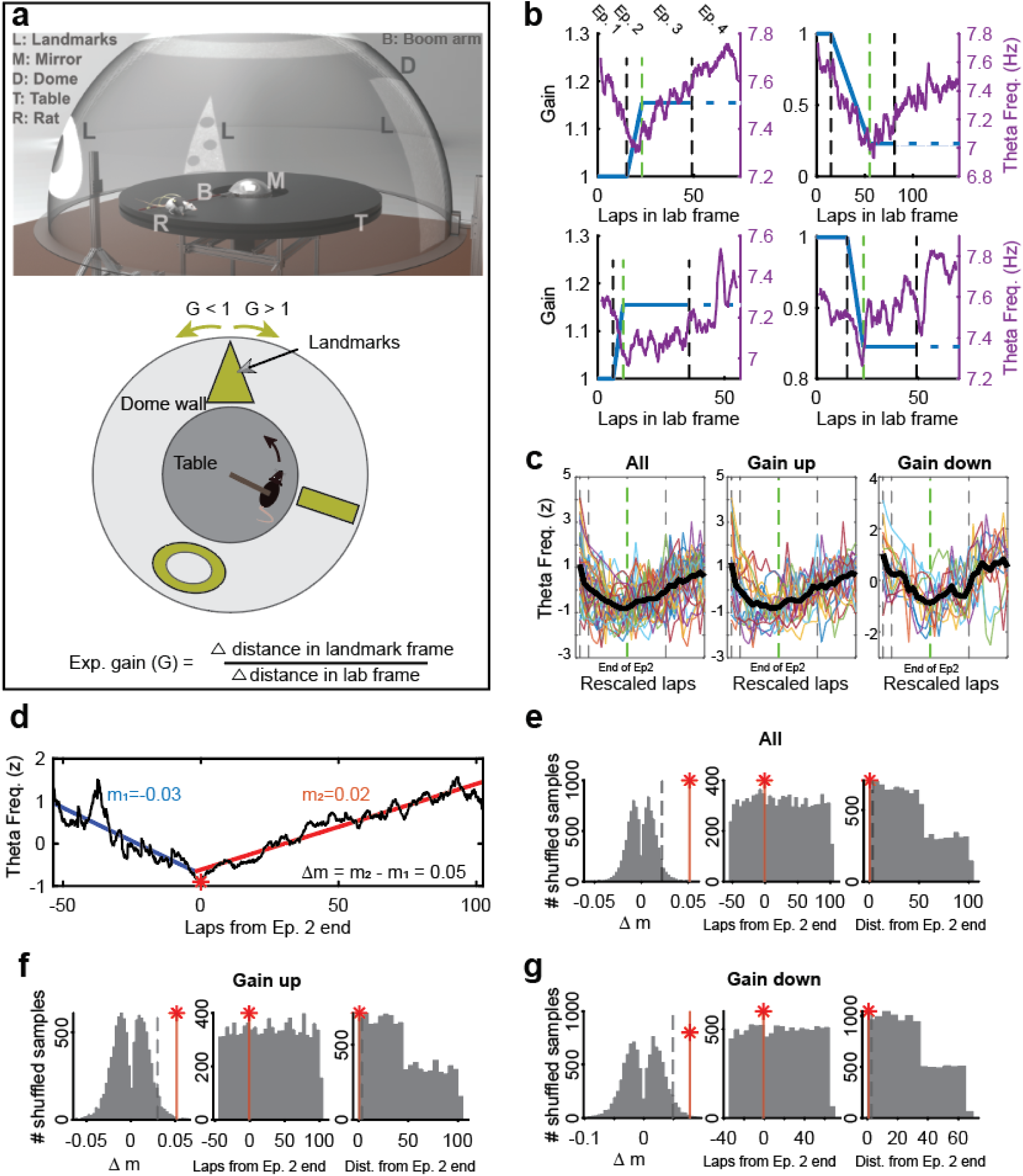
Theta frequency changes during the experiment. a. Schematic illustration of the dome environment (top) and experimental gain (bottom). *Top*: Semi-transparent illustration of the dome environment adopted from^17^. *Bottom*: A rat was fixed to a radial arm (brown) and it moved counterclockwise (CCW). Landmarks (yellow) were projected onto the dome interior and rotated in proportion to the rat’s movement, defining the “experimental gain” (G). When *G* = 1, the landmarks were stationary, and the distance traveled in the landmark frame equaled the distance traveled in the real world. When *G* > 1, the landmarks rotated at a speed proportional to the rat’s speed but in the opposite direction, causing the visual illusion that its movement was twice as fast as it was. Conversely, when *G* < 1, the landmarks rotated proportional to the rat’s speed but in the same direction, making the rat visually perceive its movement as slower than it was. b. Example theta frequency pattern during Gain up and Gain down sessions. The x-axis indicates laps in the laboratory frame of reference. The blue line indicates the experiment gain and the purple line indicates theta frequency. Four epochs were defined: epoch 1 (constant gain, Ep. 1), epoch 2 (experimental gain ramping to a target gain, Ep. 2), epoch 3 (target gain maintained, Ep. 3), and epoch 4 (landmarks turned off, Ep. 4). Vertical lines mark epoch boundaries. The dashed blue line in epoch 4 indicates the target experimental gain for reference even though the landmarks were turned off. c. Rescaled overlay plots of theta frequency across sessions. Data were aligned and scaled to epoch boundaries (vertical dashed lines) for all sessions (left), Gain up sessions (middle), and Gain down sessions (right). Each colored trace indicates a session; the black trace is the average theta frequency. All conditions exhibit a characteristic V-shaped profile, with the global minimum frequency occurring near the transition between epoch 2 epoch 3. d. Example of slopes of average theta frequency trace. The V-shaped dynamics were quantified using a piecewise linear regression model. The acuteness of the V-shape was quantified as Δm = the slope of the second segment (m_2_) – the slope of the first segment (m_1_). All sessions were aligned to the end of epoch 2 (0 on the x-axis) before the slope measurement. The red star indicates the global minimum (minimum point) of the theta frequency trace. e. Comparison with shuffled data for all sessions. Left panel shows the distribution of the slope difference between two piecewise best-fit lines (Δm) of the shuffled data. The shuffles exhibited a bimodal distribution of Δm, consistent with the known tendency of piecewise regression models to avoid 0° and 180° when fitting a continuous function^63^. The very small subset of the shuffled data that exhibited a more pronounced V-shape than the empirical data were anomalies characterized by breakpoints located at the edges of the data range (Fig. S1b). The middle panel shows a uniform distribution of laps with the theta-frequency minimum points of the shuffled data. The data were aligned to the end of epoch 2 (0 on the x-axis), with some global minimum points having a negative value because they occurred before the end of epoch 2. Because distances cannot be negative, offsets occurring before the end of epoch 2 were converted to their absolute values (*right*). This conversion resulted in a step-like null distribution, reflecting the fewer number of laps available before epoch 2 compared to after.. Vertical red lines and red stars indicate the measured data values while gray dashed lines represent the 95^th^ percentile (Δm) or the 5^th^ percentile (Distance) of the shuffled data. f. Same as (e) for the Gain up sessions only. g. Same as (e) for the Gain down sessions only.

The experiment was divided into four epochs (Fig. 1b). In the first epoch, the landmarks remained stationary (*G* = 1) to establish a baseline measurement of place fields and theta frequency. In epoch 2, *G* was gradually increased or decreased, creating an increasing mismatch between the estimated travel distance from nonvisual self-motion cues (e.g., vestibular, motor copy, and proprioceptive cues) and that inferred from visual landmarks. When *G* reached its target final value (*G*_final_) between 0.1 and 1.8, the gain was held constant for 24 to 52 laps (epoch 3), depending on the value of *G*_final_, allowing the path-integration system to adapt to the new relationship between idiothetic and allothetic cues. At the start of epoch 4, the visual landmarks were abruptly removed, unmasking the internal path-integration gain in the absence of the dominating influence of allothetic landmarks. Our previous study demonstrated that the hippocampal gain, *H*, was typically controlled by the visual landmark gain in epochs 2-3 (i.e., *H* = *G*). When visual cues were removed, this landmark-controlled hippocampal gain persisted (although reduced in magnitude), indicating that path-integration gain was recalibrated through experience^17^.

We tested whether conflicts between *G* and the internal path-integration gain induced systematic changes in theta frequency, which would thus serve as a readout of the current state of the hidden path-integration gain. Figure 1b illustrates examples of theta frequency dynamics, in which theta frequency decreased during both *G* > 1 and *G* < 1 ramping manipulations (epoch 2) and began to rebound at the start of the stable gain period after *G* reached its *G*_final_ value (epoch 3). Rescaled overlay plots in Fig. 1c revealed a V-shaped profile of the average theta frequency across laps, with a global minimum at the end of epoch 2. The same pattern emerged when sessions were aligned directly to the endpoint of epoch 2 without scaling (Fig. S1a). The V-shaped temporal profile was consistent across both gain-up and gain-down conditions (Fig. 1c, Fig. S1a).

We applied piecewise regression to the averaged theta frequency trace (collapsed across gain-up and gain-down sessions) to estimate the slopes of its descending (*left*, *m_1_*) and ascending (*right, m_2_*) segments (Fig. 1d). We defined Δm as the difference between these slopes (*m_2_ – m_1_*), where higher positive values indicate a more pronounced V-shape, zero indicates that the two segments are parallel, and negative values indicate an inverted V-shape. To verify whether the theta frequency displayed a change in the sign of the slope at the end of epoch 2, we measured the offset between the end of epoch 2 and the lap where the minimum theta frequency was observed (red star in Fig. 1d). We compared these measured values against null distributions generated from 10,000 shuffled datasets, where the theta frequency trace of each session was circularly shifted by a random value and averaged.

The Δm of frequency change was significantly higher in the empirical data (Δm = 0.051) than in the shuffled controls (p = 0.0026, Shuffle test, Fig. 1e, *left*). We compared the relative fit of the piecewise model against a linear model using the difference in Bayesian Information Criterion (ΔBIC). The empirical data yielded significantly more negative ΔBIC values compared to the shuffled data (p = 0.0002, Shuffle test, Fig. S1c), confirming that the observed V-shape is a statistically robust feature rather than a stochastic artifact. The lap with the minimum theta frequency in the data was significantly closer to the end of epoch 2 than chance (0.7 laps, p = 0.0080, Shuffle test, Fig. 1e, *right*). These effects were robust across both gain-up (Δm = 0.051, p = 0.0059; minimum point distance: 0.9 laps, p = 0.0109, Fig. 1f) and gain-down (Δm = 0.077, p = 0.0075; minimum point distance: 0.7 laps, p = 0.0117, Fig. 1g) sessions. In contrast to theta frequency, there was no systematic pattern of changes in theta amplitude except for a transient increase at the landmark-off event (Figure S2). These findings suggest that theta frequency is modulated by conflicts between internal and external positional cues in both gain-up and gain-down conditions, with theta frequency decreasing as a function of increasing conflict and then increasing as the conflict is resolved during recalibration of the path integration gain.

### Theta frequency change during allothetic–idiothetic cue conflict predicts path-integration gain recalibration

We next examined whether changes in theta frequency during gain manipulation could predict the magnitude of hippocampal path-integration gain recalibration. We computed the difference in hippocampal gain (*dH*) right before (*H*_final_) and right after (*H*_recal_) the landmarks were turned off (Fig. 2a). *H*_final_ reflects the last hippocampal gain controlled by the visual landmarks and *H*_recal_ represents the first hippocampal gain measured in the absence of landmark influence, which we interpret to reflect primarily the path-integration gain.

**Figure 2.**
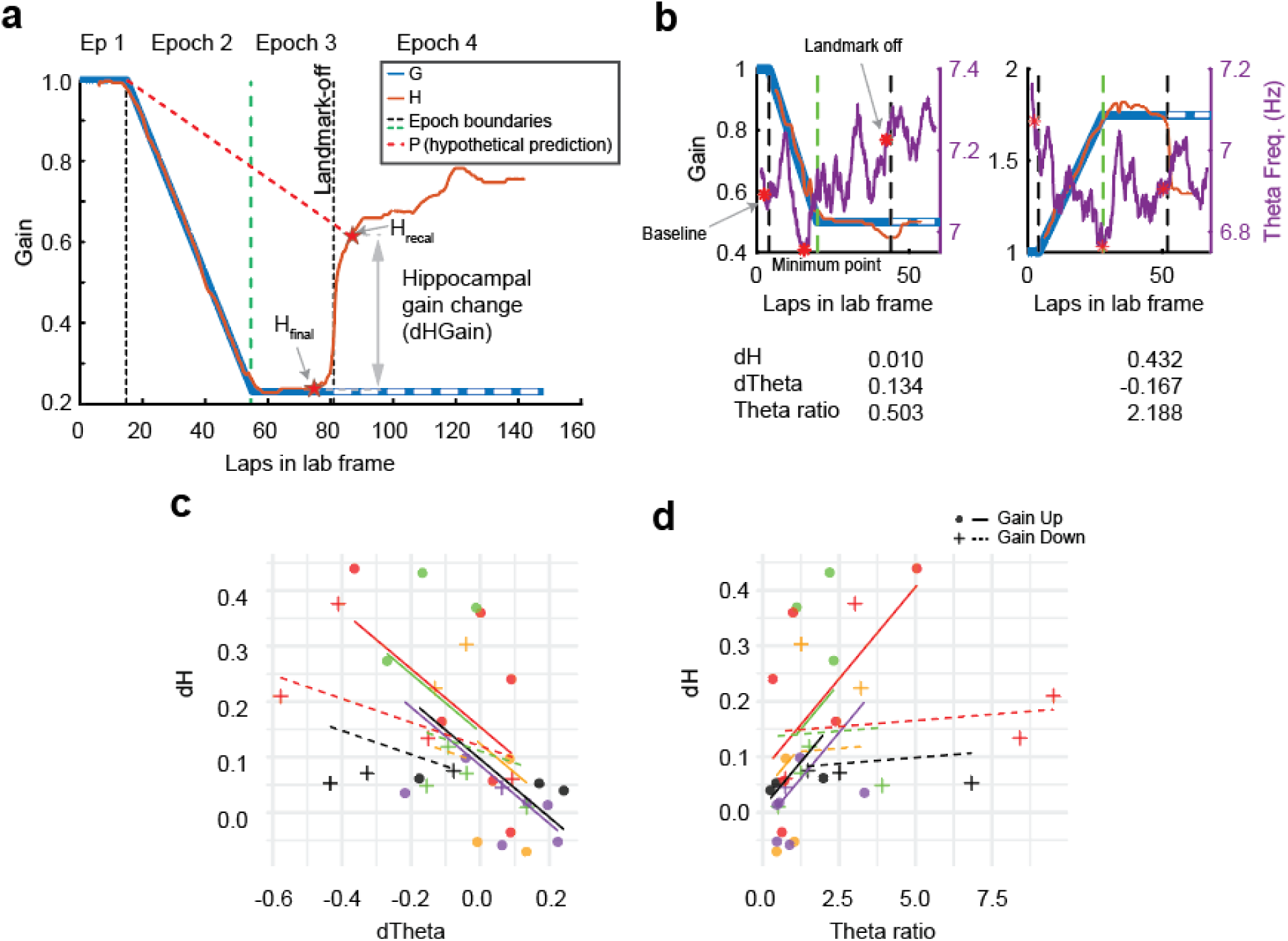
Theta frequency dynamics predict hippocampal gain adjustments following landmark removal. a. Method for calculating the change in hippocampal gain (dH) following landmark removal. Blue and orange lines indicate the experiment gain (G) and hippocampal gain (H), respectively. dH is the difference between the final landmark-driven hippocampal gain (point ‘H_final_’, measured at the center of the last 12-lap analytical window prior to landmark removal) and the initial path-integration-driven hippocampal gain (point ‘H_recal_’, measured at the center of the first 12-lap analytical window after the landmark removal). Specifically, we chose the convention *dH* = *DIR* × (*H*_final_ − *H*_recal_), where *DIR* is a directional indicator, defined as +1 for gain-up sessions and −1 for gain-down sessions. In this convention, *dH* > 0 reflects incomplete recalibration, as the path-integration gain was at a value somewhere between the epoch 1 hippocampal gain (≈ 1) and *H*_final_, regardless of the direction of the landmark gain change (*dH* > 0 could also indicate “negative” calibration, e.g., when *H*_final_ is 0.8 and *H*_recal_ is 1.1, although such cases were rare.) A *dH* of zero indicates that the path-integration gain measured after the landmarks were turned off was identical to the hippocampal gain just before the landmarks were turned off, demonstrating complete recalibration of the path-integration gain. *dH* < 0 indicates over-calibration which happened on a small proportion of sessions. The red dashed line (P) illustrates the hypothesized, unobservable state of the internal path-integration gain during the presence of the landmarks. b. Representative single-session examples. The panel displays hippocampal gain as in panel (a) alongside the corresponding theta frequency (purple trace). The calculated values for dH at the landmark removal event, dTheta (the difference in theta frequency between a 3-lap average at the end of epoch 1 and a 3-lap average at the end of epoch 3), and the theta ratio for these sessions are shown below each plot. Critical reference points for dTheta and theta ratio (baseline, minimum point, and landmark-off) are indicated by red stars. c. A linear mixed-effects model, with the direction of gain manipulation (gain-up and gain-down) as an additional fixed effect and individual animal as a random effect, showed that dTheta was negatively correlated with dH. (The random effect of subjects allowed random intercepts only, to allow the model to converge; see Fig. S3a-b for the full random effect model and the random-slope-only model.) Markers indicate gain direction (fixed effect) and colors denote individual animal identity (random effect); dashed and solid lines represent the regression fits for Gain down and Gain up conditions, respectively. d. Correlation between theta ratio and hippocampal gain change. A linear mixed-effects model revealed a significant positive correlation between the theta ratio and dH.^64^ The same format is used as in panel c.

Our goal was to test whether theta frequency dynamics reflect the magnitude of the mismatch between *H*_final_ and the hidden path-integration gain at the end of epoch 3 and thereby predict the magnitude of path-integration gain recalibration revealed in epoch 4. We derived two independent variables based on theta frequency trajectories (Fig. 2b). (1) *dTheta* was defined as the difference in theta frequency between the end of epoch 1 (*baseline*) and the end of epoch 3 (*landmark-off* event). (2) *Theta ratio* was defined as the decrease in theta frequency from the *baseline* to the global minimum (*minimum point*) divided by the increase from the *minimum point* to the *landmark-off* event. A lower theta ratio indicates a stronger recovery (typically during epoch 3) of theta frequency after the initial drop (typically during epoch 2).

We first investigated the relationship between *dTheta* and *dH* using a linear mixed-effects model^26^. Greater reductions in theta frequency by the time of landmark removal were associated with greater deviations between *H*_final_ and *H*_recal_ (β = –0.524, 95% confidence interval (CI) = [-0.852,-0.199], p < 0.05, Bayesian Information Criterion (BIC) = –15.6; Fig. 2c). This result shows that the net change in theta frequency during the experimental gain manipulation predicts the mismatch between the final hippocampal gain and the recalibrated path-integration gain.

We next investigated the relationship between the *theta ratio* and *dH* to determine whether the proportionate amount of rebound of theta frequency after its initial decrease predicted the magnitude of path-integration gain recalibration. The model revealed that the decreased *theta ratio* was associated with decreased *dH* (β = 0.066, CI = [0.019,0.116], p < 0.05, BIC = –3.4; Fig. 2d, S3c-d), indicating that a higher recovery of theta frequency is associated with less error between the hippocampal and path-integration gains and a greater degree of recalibration. Together, these results show that both the cumulative change in theta frequency (*dTheta*) and the shape of its trajectory (*theta ratio*) reflect the degree of mismatch between path-integration gain and hippocampal gain at the point of landmark removal. This relationship was not an artifact of speed– or acceleration-based modulation of hippocampal theta (Fig. S4)^24, 25, 27–29^.

### Theta frequency as a continuous readout of the error between path-integration gain and hippocampal gain in individual sessions

The prior analyses were based on the theta frequencies averaged over many sessions. We next investigated whether the theta frequency change during epochs 2 and 3 could reflect the moment-to-moment magnitude of error between hippocampal gain and path-integration gain across laps during the dynamic recalibration process itself. Although path-integration gain cannot be directly measured while the landmarks are on, we inferred its temporal dynamics based on 3 assumptions:

1) Path-integration gain in epoch 1 (with stationary landmarks) is approximately equal to the hippocampal gain (i.e., Path-integration gain ≈ 1).
2) Path-integration gain is unaffected by the landmark removal event itself and is approximately the same immediately before and after the landmarks are turned off.
3) Path-integration gain changes approximately linearly while the landmarks are rotating according to the current value of *G* in epochs 2 and 3.

Based on assumptions 1 and 2, we estimated the initial and final values of path-integration gain during the landmark-on period as, respectively, the values of *H* at the end of epoch 1 and at the 6th lap (center of the first 12-lap window) after the landmarks were turned off. Under assumption 3, we linearly interpolated path-integration gain between these two points to estimate its lap-by-lap changes, a simplified estimate that assumes that path-integration gain changes gradually and continuously (Fig. 3a). (An alternative approach using a Kalman filter to infer the lap-by-lap hidden path-integration gain produced similar results [not shown]).

**Figure 3.**
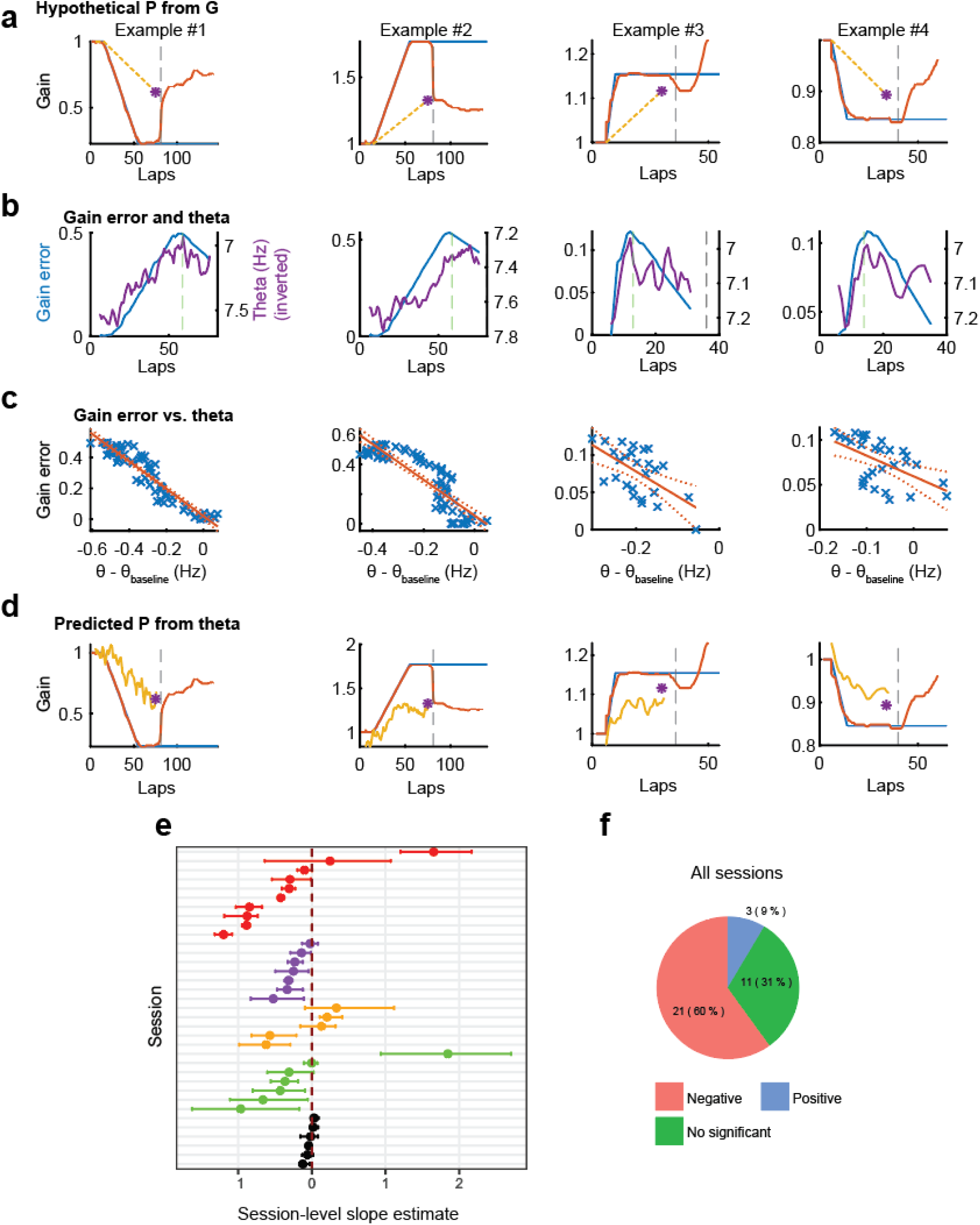
Lap-by-lap estimation of gain error from theta frequency. a-d. The analysis pipeline is illustrated using four representative sessions (see also Fig. S5). Each column presents data from a single session, while rows represent the sequential steps of the analysis pipeline. a. Estimation of the lap-by-lap hypothetical path-integration gain (*P*, yellow dashed line) via linear interpolation between the end of epoch 1 and the pre-landmark-off point. The blue and orange lines represent the experimental gain (*G*) and hippocampus gain (*H*), respectively. The gray dashed line represents the landmark-off event. b. Lap-by-lap gain error and theta frequency. The gain error (blue line), defined as the difference between hippocampal gain (*H*) and the estimated path-integration gain (*P*) from panel (a), is plotted alongside the theta frequency (purple line), which is shown on an inverted y axis to align with the H values for visual comparison. The green dashed line represents the end of epoch 2. c. Linear regression of gain error against theta frequency. A linear model was fit to predict gain error from the deviation of theta frequency from its baseline (θ – θ_baseline_). Blue crosses represent individual laps, and the orange line indicates the linear fit with 95% confidence bounds. d. Reconstructed path-integration gain. To evaluate the model’s predictive power, the path-integration gain (*P*, yellow line) was reconstructed based on theta frequency using the linear relationship established in panel (c). e. Summary of regression slopes across all analyzed sessions. The plot displays the estimated slopes and 95% confidence intervals (determined through a bootstrap procedure; see Methods) for the linear models fit to each session. Colors denote individual animal identity. f. Classification of sessions by model significance. The pie chart categorizes all sessions based on whether the regression slope was significantly negative (red), significantly positive (blue), or not significant (green).

We estimated lap-by-lap gain error magnitude (the difference between the linearly interpolated path-integration gain and the measured hippocampal gain) and examined its similarity to the theta-frequency change (Fig. 3b; we inverted the y axis of the theta frequency to align the two plots). Notably, at the end of epoch 2, where the gain error peaked, the theta frequency typically reached a minimum, as gain error and reversed theta frequency showed similar trajectories (Fig. 3b, S5). Using a linear model, we tested whether gain error could be predicted from theta frequency change on a lap-by-lap basis (Fig. 3c) as below:

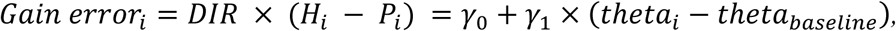

where *Gain error*_*i*_ indicates the gain difference between interpolated path-integration gain (*P*_*i*_) and hippocampal gain (*H*_*i*_) in lap *i*; (*theta*_*i*_ − *theta*_*baseline*_) indicates the difference between theta frequency in lap *i* and the theta frequency at the end of epoch 1; and *DIR* is a directional indicator, defined as +1 for gain-up sessions and −1 for gain-down sessions. Representative examples showed significantly negative slopes, indicating that theta frequency change can predict gain error (Fig. 3c). We then reconstructed the path-integration gain trajectory based on this relationship between gain error and theta frequency:

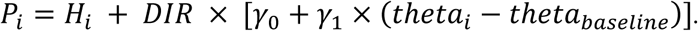

The resulting estimate closely matched the linearly interpolated path-integration gain inferred from the hippocampal gains at the beginning and end of the landmark-on period (Fig. 3d).

We observed that 24 out of 35 sessions (69%) demonstrated a significant correlation between gain error and theta frequency (p < 2.2 × 10^-16^, binomial test against a chance percentage of 0.05, Fig. 3e-f). Of these 24 sessions, a significantly larger portion (21 sessions) showed a negative slope than a positive slope, consistent with the population-level analysis in Fig. 2c (binomial test, p = 0.0003 compared to the null hypothesis of equal numbers of positive and negative correlations). While acknowledging that the precise measurement of path-integration gain and gain error cannot be directly achieved and relies on estimations, these findings nevertheless provide compelling evidence that theta frequency correlates with lap-by-lap gain error.

### Theta frequency predicts gain mismatch in landmark failure sessions

In some sessions, the hippocampal gain began to deviate from the experimental gain when the experiment gain was decreasing in epoch 2^17^; that is, the place cell map began to drift relative to the landmark frame of reference. We analyzed whether theta frequency retains the predictive power over gain recalibration in these “landmark-failure” sessions. We first examined individual sessions and observed that the decreasing trend in theta frequency often ceased around the landmark-failure point, that is, the point when the hippocampal gain and landmark gain diverged (Fig. 4a). Unlike in landmark-controlled sessions, the average theta frequency trace did not exhibit a clear V-shape with its minimum at the end of epoch 2. Instead, the initial decrease in theta frequency halted at the landmark-failure point in epoch 2 and was followed by a period of fluctuation around a relatively flat mean before rising again around the landmark-off event (Fig. 4b; S6a-d).

**Figure 4.**
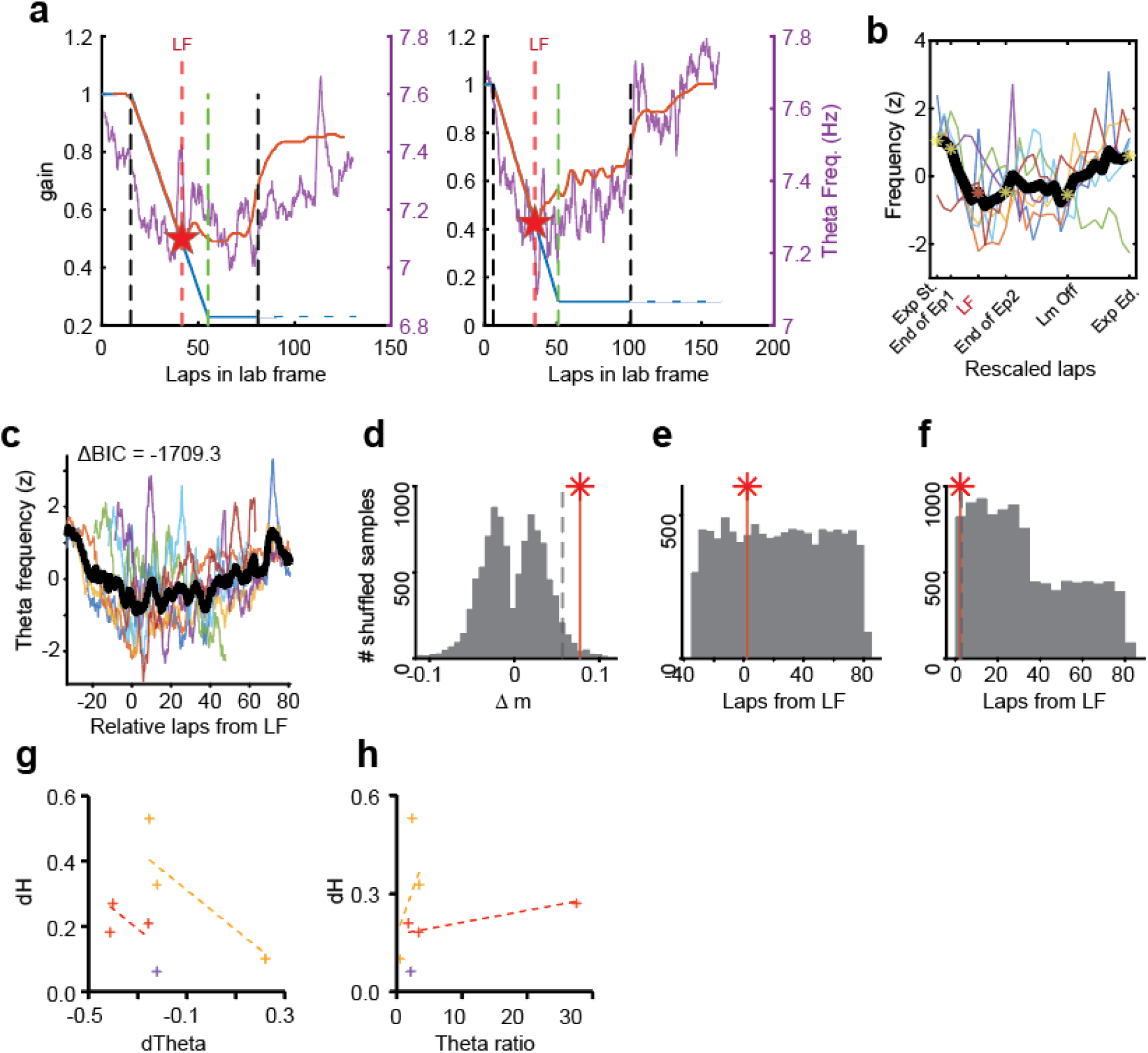
Theta frequency as a potential predictor of gain mismatch during landmark-failure sessions at the population level. a. Representative session examples illustrating landmark-failure (LF) events (i.e., when the hippocampal gain diverges from the experimental gain, marked by red stars and vertical lines) occurring during epoch 2 of Gain down conditions. The experimental gain (G, blue), hippocampal gain (H, orange), and theta frequency (purple) are plotted. b. Realigned overlay of theta frequency traces across landmark-failure sessions. All sessions (colored traces) and their average (bold black trace) are shown with epochs individually rescaled to align epoch boundaries and the LF event. In contrast to landmark-controlled sessions, the average theta frequency leveled off after the LF event. c. Overlay of theta frequency traces aligned by the landmark-failure (LF) event. ΔBIC values are provided to compare the piecewise and linear regression fits for data aligned to the LF point. Aligning the data to the landmark-failure point produced a ΔBIC of –1709.3, whereas aligning to the end of epoch 2 yielded a ΔBIC of –875.8 (Fig. S6a). This ΔBIC difference (–833.5) indicates that the theta-frequency dynamics are more systematically structured around the landmark-failure point than around the end of epoch 2^65, 66^. d. Shuffle test for the Δm of the V-shaped theta frequency profile, where overlay plot is aligned to the landmark-failure event. The observed value from the group average is marked with a red star and a vertical line, with the histogram representing the distribution of the shuffled data. The gray dashed line represents the 95th percentile of shuffled data. e. Distribution of the lap with the theta frequency global minimum from shuffled data. The x-axis indicates relative laps from the end of epoch 2; the observed value from the group average is marked with a red star and a vertical line. f. Shuffle test for the distance of the lap with the theta frequency global minimum of the V-shaped theta frequency profile to the landmark failure point. The gray dashed line represents the 95th percentile of shuffled data. g. Relationship between dTheta and dH, fit with a linear mixed-effects model with rat identification as a random effect. The direction of landmark gain change was not used as an additional fixed effect because all landmark-failure sessions were gain-down sessions^17^. Dotted lines indicate the linear fit, and colors represent individual animals. A random-intercept-only model was utilized for this analysis. h. Relationship between the theta ratio and dH, fit with a linear mixed-effects model. A random-slope model was applied instead of a random-intercept model, as the model resulted in a singular fit.

When we realigned the data to the landmark-failure point (*LF*) of each session (Fig. 4c), the *Δm* of the V-shape became significantly more acute than chance (Δm = 0.077, p = 0.0185, shuffle test, Fig. 4d) and the lap with the global minimum of theta frequency was a significantly shorter distance from the landmark-failure point than predicted by chance (2.3 laps, p = 0.0394, shuffle test, Fig. 4e and f). We also tested whether *dTheta* or *theta ratio* predicted *dH* in the landmark-failure sessions.^17^ While *dTheta* significantly predicted *dH* (β = –0.608, CI = [-1.049,-0.135], p < 0.05, BIC = 2.5, linear-mixed model with random intercept, Fig. 4g, Fig. S6e), *theta ratio* did not, likely due to the small number of landmark-failure sessions minimizing the statistical power (β = 0.018, CI = [-0.059,0.100], p > 0.05, BIC = 12, linear-mixed model with random slope, Fig. 4h, Fig. S6f). Taken together, these results show that the modulation of theta frequency was highly related to the dynamics of hippocampal gain even in landmark-failure sessions.

Analysis of individual sessions revealed that theta frequency can significantly predict the error between path-integration gain and hippocampal gain in landmark-failure sessions. Out of 7 sessions, theta frequency significantly predicted gain error in 5 sessions (71.4 %, p = 6.027 × 10^-06^, binomial test against a chance percentage of 0.05, Fig. 5a-c). Furthermore, in 4 of the 5 sessions with significant prediction, the coefficient of theta frequency was significantly negative. Although this proportion is not significant based on a 2-tailed binomial test, likely due to the small sample size of landmark-failure sessions (80.0%, p = 0.3750, compared to the null hypothesis of equal numbers of positive and negative correlations, Fig. 5c), it is similar to the pattern observed in the landmark-controlled sessions (Fig. 3f). The session-wise analyses suggest that even in landmark-failure conditions, theta frequency serves as a useful predictor of gain error in most sessions.

**Figure 5.**
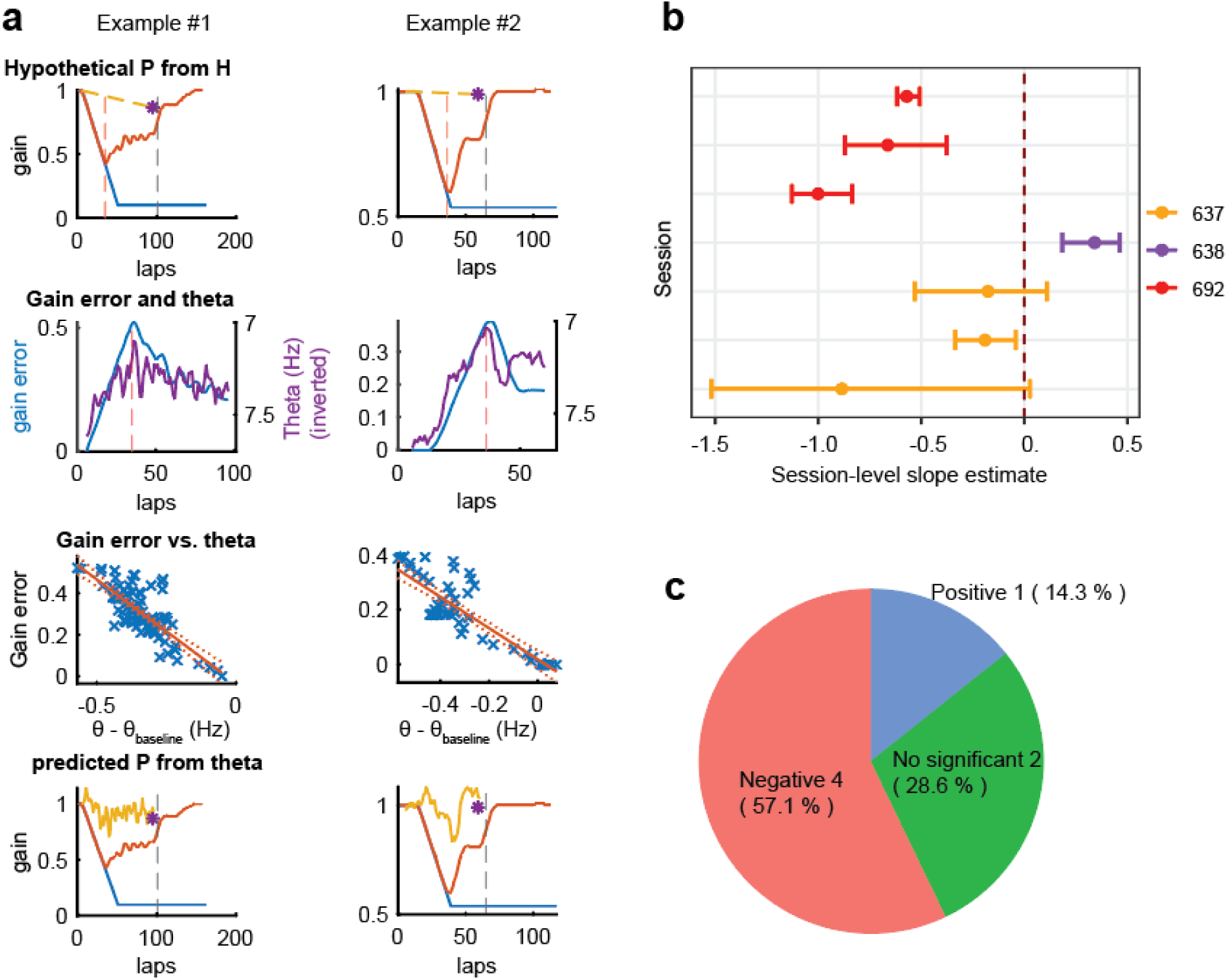
Theta frequency predicts lap-by-lap gain error at the individual session level. a. Illustration of the single-session analysis pipeline using representative examples of landmark-failure sessions. Each row depicts a sequential step in the analysis. See Figure 3a-d for details. b. Summary of linear model slopes across all landmark-failure sessions, displaying the slope estimate and 95% confidence interval for each session. c. Session classification by model significance. The pie chart categorizes the sessions from (b) based on whether the slope was significantly negative (red), significantly positive (blue), or not significant (green).

### Continuous attractor neural network model predicts theta frequency decreases by gain discrepancy

A key result of these experiments is that the theta frequency decreases regardless of the sign of the error between the path integration gain and the hippocampal gain (i.e., in both *G* > 1 and *G* < 1 sessions), rather than changing linearly with *G* (i.e., decreasing frequency as *G* decreases below 1 and increasing frequency as *G* increases above 1). To gain insight into the network dynamics and mechanisms that might produce such a symmetric response to the positive and negative gain errors, we investigated a recent continuous attractor neural network (CANN) model (Fig. 6a) developed by Chu et al.^30^, which our group previously modified to gain insight into the dynamics of theta phase precession during cue-conflict and gain recalibration experiments^31^.

**Figure 6.**
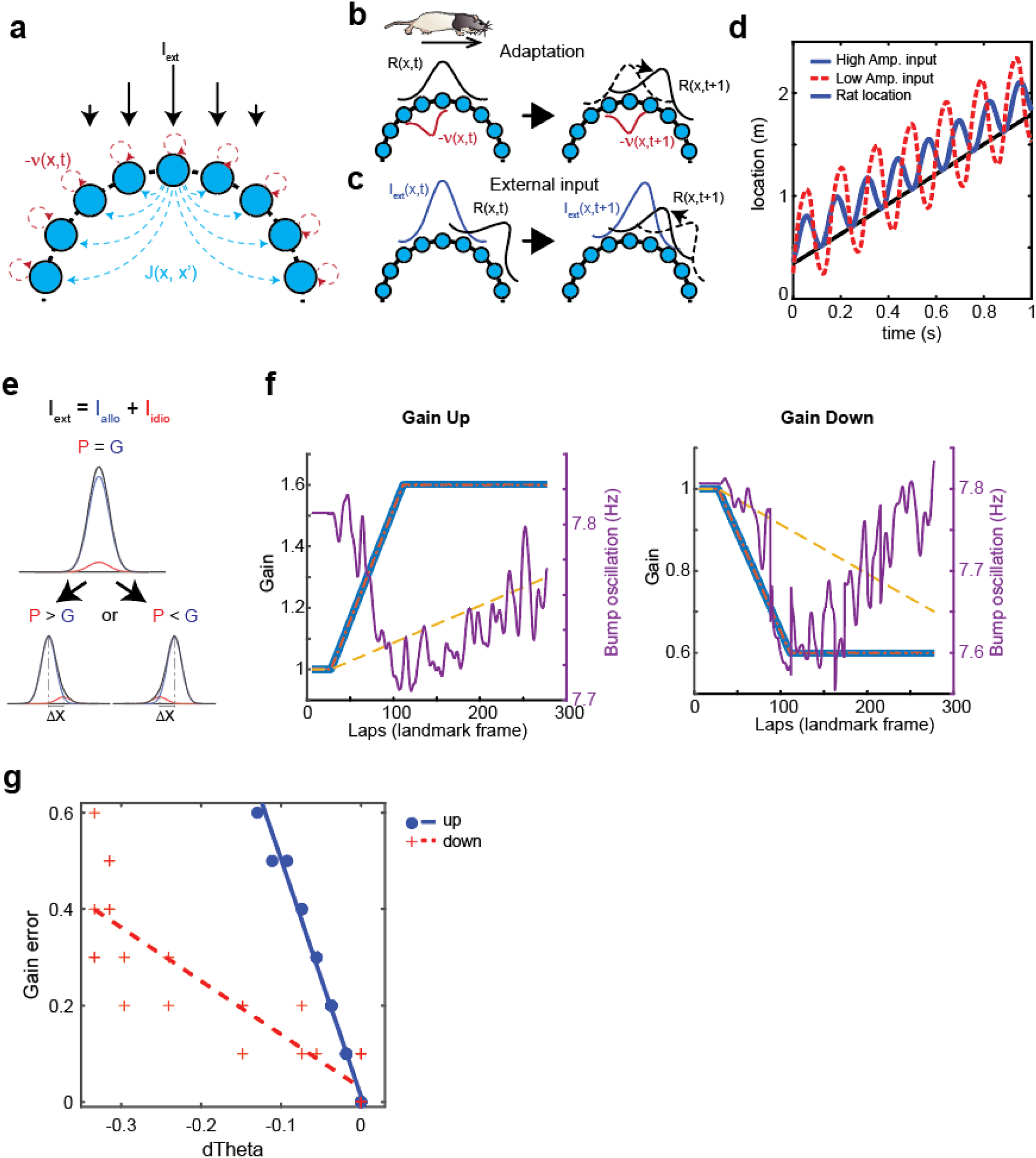
Continuous Attractor Neural Network (CANN) model predicts theta frequency attenuation induced by gain discrepancy in both directions. a. Schematic representation of CANN model, adapted from Reference ^30^. Neurons (blue circles) are arranged in a ring representing location on a circular track (*x*). Synaptic drive in this model is governed by three key factors: recurrent connections (*J*), firing rate adaptation (*v*), and an external input (*I*). b. Effect of adaptation on the activity bump in a moving rat. Schematic illustration of neural activity as a rat runs clockwise on a circular track. The left panel shows the population firing pattern (’bump’) of place cells (blue circles) representing continuous space (*x*) at a specific time (*t*). Adaptation (−*v*) acts similarly to an inhibitory input generated by prior synaptic activity. Because it is influenced by past synaptic inputs, the adaptation is misaligned with the current bump in a moving rat. This imbalance destabilizes the bump on the attractor ring and serves as a primary driver for its movement away from the neurons undergoing adaptation (right panel). (Rat image adapted from Scidraw.io ^67^) c. Schematic illustrations of attractor modulation by external input (*I*). Blue and black lines represent the external input and the activity bump on the attractor ring, respectively. In contrast to adaptation, the external, excitatory input exerts an attractive force pulling the bump toward it (right panel). Together with adaptation, the external input serves as a critical driver for the attractor’s oscillation. d. Oscillation of the continuous attractor as a function of input strength. Comparison of oscillations in the same network under high amplitude input (blue solid line, *alpha* = 0.11) and low amplitude input (red dotted line, *alpha* = 0.08). The result shows a 1-second simulation of bump oscillation in a rat moving forward. The x-axis represents time, and the y-axis represents the bump’s position on the ring. While both conditions share a similar phase during the first cycle, the high-input condition (blue) oscillates over a narrower range of positions and at a faster frequency than the low-input condition (High amplitude input: 7.82 Hz; Low amplitude input: 6.82 Hz). The black line shows the rat’s spatial position over time. e. Input amplitude modulation via spatial misalignment. This panel Illustrates how the spatial alignment between allothetic (G, blue) and idiothetic (P, orange) inputs affects the resultant amplitude of the total input (black). *(upper)* When inputs are aligned (G = P = 1.0), the summed excitatory drive is maximized. *(lower)* A gain mismatch creates a spatial offset between the two inputs, reducing the peak amplitude of total excitation. f. Representative simulation example for Gain up *(left)* and Gain down *(right)* sessions. Epoch 1 to epoch 3 are simulated. These panels show the evolution of hippocampal gain (orange) and the theta-frequency oscillation of the neural activity bump (purple line) as the experiment gain (blue) and path-integration gain (yellow dashed line) are changed. The hippocampal gain closely tracked the experimental gain. g. Correlation between gain error and theta frequency shift. Simulation results illustrate the predicted change in gain error, defined as the difference between the hippocampal gain and the path integration gain, as a function of the change in theta frequency (dTheta). The blue solid line depicts the prediction for the Gain up condition, while the red dashed line represents the prediction for the Gain down condition. In addition to the significant main effect of *dTheta*, the linear model also reported a significant interaction between *dTheta* and gain direction (β = 3.812, CI = [3.160,4.463], p < 0.05; Fig. 6g). This effect is primarily due to an asymmetry between Gain up and Gain down sessions in the scale of the gain values (i.e., gain up magnitudes can range from 1 to infinity in principle, whereas gain down magnitudes are limited from 0 to 1).

In this model, the authors explain that two interacting forces drive the activity bump of a ring attractor to oscillate between two positions near the rat’s actual position^30^. Adaptation, acting similarly to an inhibitory input based on prior synaptic activity, provides inhibition to the area corresponding to the previous bump position in a moving rat (i.e., the position just behind the rat). This adaptation makes the bump unstable, allowing it to move away from its current position (Fig. 6b). Conversely, external input acts as a pulling force that brings the bump back toward the rat’s position (Fig. 6c). Stronger input counteracts the repulsive drive of the adaptation and arrests the forward movement of the bump more quickly than weaker input, shortening the bump’s length of travel along the ring and increasing the frequency of the oscillation (Fig. 6d).

Following Sueoka et al.^31^, we modified the Chu et al.^30^ model by introducing two input streams: allocentric input (reflecting landmark gain) and idiothetic input (reflecting path-integration gain; Fig. 6e). These gains were manipulated analogously to experimental procedures (landmark gain ranged from 0.3 – 1.7, while path integration gain was constrained between the landmark gain and 1). In this simulation context, we examined the relationship between the *Gain error* (defined as the difference between the hippocampal gain measured from hippocampal bump and the simulated path integration gain) and the change in theta frequency (*dTheta*, defined as the change in theta frequency relative to the baseline condition where both gains equaled 1), using the same analyses performed on the empirical data (Fig 2c and 3a-c). Figure 6f demonstrates the simulated changes in bump-oscillation frequency and hippocampal gain under the landmark-gain and path-integration-gain manipulations for gain-up and gain-down sessions. Regardless of the direction of the gain change, theta frequency in epoch 2 decreased as the discrepancy between the landmark gain and the path-integration gain increased, and in epoch 3 it increased as this discrepancy decreased, just as observed in the empirical data. A linear model analysis, using the gain direction and *dTheta* as independent variables, revealed a reduction in *Gain error* as *dTheta* increased (β = –4.921, CI = [-5.400,-4.603], p < 0.05; Fig. 6g, BIC = –143.4;Fig. 6g, Table S1), as we observed in the recording data in Fig. 2c.

## Discussion

A primary challenge in understanding the neural bases of cognition is the influence of hidden internal factors that are difficult to measure directly^1, 2^. Path integration, which is essential for building and updating cognitive maps, is an example of such a hidden computation under conditions in which its operation is masked by the overriding influences of landmark navigation processes^6, 15, 16^. In this study, we discovered a novel method to indirectly estimate path-integration gain even in the presence of dominating allothetic inputs by measuring the relationship between the observable hippocampal gain and the modulation of theta frequency. Theta frequency decreased continuously during the period in which the experiment gain (*G*) was ramping up or down (epoch 2) but began to increase when the ramping stopped (epoch 3). During these epochs, the hippocampal gain was typically driven by visual landmarks. In contrast, after the landmarks were turned off, hippocampal coding was liberated from the influence of visual landmarks, revealing the recalibrated path-integration gain^17^. We found that both the accumulated change in theta frequency (*dTheta*) *and* the pattern of its change (*theta ratio*) predicted the error between the landmark-controlled hippocampal gain and the path-integration gain (*dH*) around the landmark-off event: greater net reduction and less-complete recovery of theta frequency were associated with less recalibration of hippocampal gain (Figure 2c,d). In landmark-failure sessions, theta frequency dynamics shifted, with changes occurring near the point of landmark failure rather than at the end of epoch 2.

### Beyond direct representation: a novel way to estimate hidden variables indirectly

Beyond the specific domain of spatial navigation, our findings provide a conceptual framework for how hidden cognitive variables or computations might be estimated. In many cases, electrophysiology and imaging enable direct observation of how cognitive processes are implemented through physiological activity. Prime examples are the discovery of place cells in the hippocampus, which form an allocentric representation of an animal’s location^32^, and head-direction cells, which directly represents the animal’s current allocentric head direction^33^. The common thread in these studies is that the physiological signal appears to directly represent the cognitive variable of interest.

However, this approach can fail when it is unclear whether the variable is explicitly represented in neural spiking activity, where in the system the variable is explicitly computed, and whether its representation can be separable from representations of other variables in the system. In the case of navigation, the process of path integration is intertwined with the process of landmark-based navigation. Although the medial entorhinal cortex is thought to be a hub of path-integration computation^34–36^, even at this level it is difficult to tease apart the influences of landmarks and path integration when both are present.

In most models of path integration of head direction cells, grid cells, and place cells, the path integration gain is implicitly represented in the distribution of synaptic weights from representations of speed and direction to the representations of position and other properties of the network^35, 37, 38, 39 Secer, 2024#13153^. It thus may be challenging, when landmarks are available, to directly observe the specific contribution of path integration by measuring neural output *in vivo*. One could in principle measure the synaptic weights and excitatory potentials, but that is difficult to do with current technologies and would require knowing which synapses carry the path integration gain.

Our approach, in contrast, was to identify a continuously modulated neural signal (theta frequency) that serves as a proxy for an error signal (gain discrepancy) and to estimate the unobservable hidden variable (path-integration gain) with an observable variable (*H*). This methodology shares conceptual similarities with approaches in neuroeconomics, where the subjective value of a stimulus is inferred from the relationship between delivered rewards and the resulting prediction error signals by dopamine activity^40, 41^. However, a key distinction lies in the temporal nature of the estimation. While error estimation typically relies on discrete signals time-locked to specific events like reward delivery or cue onset, our study demonstrates that hidden variables can be estimated continuously and quantitatively in real-time, even in the absence of explicit reward feedback.

### What drives the change in theta frequency?

Multiple factors affect hippocampal theta frequency. For example, numerous studies in both humans and rodents reporting a positive correlation between theta frequency and movement speed^24, 25, 29, 42, 43^ or acceleration^28^ [but see ^29^]. In the present study, we confirmed that locomotion-adjusted theta frequency predicted the gain error, demonstrating that the relationship between gain error and theta frequency cannot be explained by speed or acceleration alone (Fig. S4). Another well-documented modulator of theta frequency is environmental novelty^24, 25, 44^. While these factors can explain the qualitative direction of theta frequency changes, they do not fully explain how these changes can quantitatively represent gain error. We propose a possible mechanism in which a continuous attractor neural network model can explain this relationship. The model of Chu and colleagues^30^ shows that a decrease in the amplitude of input signals will lead to a decrease in theta oscillation frequency. In our experiment, we conceptualize the inputs as two distinct representations of location: one derived from allothetic cues and the other from idiothetic cues^31^. A spatial discrepancy between these inputs induces a decrease of amplitude in the summed input, resulting in a reduction in theta frequency. This model demonstrates a possible mechanism that a gain error between *H* gain and path-integration gain can quantitatively account for the observed decrease in theta frequency, regardless of direction.

While the theta frequency was measured from the LFP in our experiments, the theta frequency in the model specifically refers to the oscillation of a population activity “bump” generated by theta sweeps of place cells^30^. Because LFPs reflect only partially the spiking activity of cells and are thought to reflect primarily synaptic currents, it is not clear how strongly this bump oscillation can be reflected in LFP theta frequency^45^. One hypothesis is that the bump oscillations originate in CA3 and subsequently influence the CA1 LFP theta frequency by virtue of the strong inputs from CA3 via the Schaffer collaterals^46^. If this hypothesis is correct, we would predict that the spiking activity of the CA3 place cells during the same experiment would show a progressively slowing oscillation during epoch 2 and a progressively speeding oscillation during epoch 3, just as we saw in the LFP signals of CA1. Another non-exclusive possibility is that the observed changes in CA1 LFP theta frequency are not a local intrahippocampal dynamic but rather emerge from a larger network-level interaction involving the hippocampus-medial septum feedback loop. While the hippocampal theta rhythm is modulated by the MS, MS cells are also regulated by direct and indirect feedback from the hippocampus^47–51^. This suggests that even if bump oscillations do not directly alter the local LFP theta frequency within the hippocampus, they could indirectly influence CA1 LFP theta frequency by modulating the MS via this feedback loop. If this hypothesis is correct, we would expect to observe consistent changes in theta frequency not only in CA1 but also in other regions modulated by the MS, such as CA3 and entorhinal cortex. Although further research is needed to elucidate the precise mechanism of theta frequency change (and there might be more than one mechanism), our model remains valuable as it provides a quantitative explanation for how theta frequency can encode the magnitude of gain error.

### Functional contribution of theta frequency modulation to the recalibration of path integration

Our data reveal a correlation between changes in theta frequency and the discrepancy between the two internal gains, but a causal link cannot be established from these results alone. To explore this question, we must consider the process of path-integration recalibration itself. According to the work of Secer and colleagues^39^, one possible mechanism for recalibration is the modification of synaptic weights in the speed-processing input into an attractor network. For example, in response to a gain increase (gain > 1) in a rat moving counter-clockwise (CCW), the synaptic weights between speed-encoding cells and the CCW rotation ring are strengthened, while the weights onto the CW rotation ring are weakened. This allows the activity “bump” in the central ring to move faster for the same physical running speed. Conversely, in response to a gain decrease, the weights onto the CW ring are strengthened and those onto the CCW ring are weakened, slowing the bump’s movement.

A correlation between reduced theta frequency and enhanced synaptic plasticity is indirectly supported by studies of working memory in humans, as recent studies reported that working memory is accompanied by synaptic plasticity^52, 53^. In one study, theta frequency in the hippocampus decreased as memory load increased^54^. In another study, during high memory load, the power of the low-theta band (4-5 Hz) significantly increased relative to the alpha band (10-12 Hz) in parts of the medial temporal lobe (MTL), including the posterior hippocampus^55^. To the best of our knowledge, there is at present no direct evidence that a reduction in theta frequency causes an increase in synaptic plasticity. However, if such a relationship exists, a lower theta frequency might contribute by extending the temporal window in which plasticity is available. For example, it has been reported that when measuring synaptic potentiation at the Schaffer collateral-CA1 synapse relative to the theta phase in the hippocampal fissure, depotentiation is favored at the peak of theta, while potentiation is favored at the trough^56–59^. Considering that a decrease in theta frequency lengthens the duration of each cycle, this extends the preferred temporal window during which a synapse is in a state permissive for either potentiation or depotentiation, thereby increasing the probability that synaptic plasticity can occur.

In conclusion, we propose a novel method to indirectly estimate path-integration gain, which is a variable typically unobservable during landmark availability, by using theta frequency and hippocampal gain. Given a computational model suggesting that gain may be encoded at the level of synaptic weights, our findings imply a potential link between the observed reduction in theta frequency and synaptic plasticity^39^. This interpretation indirectly supports Hasselmo’s SPEAR model, which posits that distinct phases within a theta cycle preferentially facilitate either synaptic potentiation or depotentiation^58, 60^. Considering that a lowered theta frequency elongates the duration of each phase within a cycle, we propose that this extends the temporal window for phase-specific plasticity, thereby increasing the probability of its occurrence. However, it should be noted that our current results do not establish a causal relationship between reduced theta frequency and enhanced synaptic plasticity. Further investigation is required to elucidate this causality.

## Materials and Methods

This study is an analysis of data previously published in several reports for different purposes^17, 18, 31^. While full methodological details on data collection can be found in the original publications, the relevant sections are briefly summarized below. In addition, all details of the new analyses are included below.

### Subjects

Five Long-Evans male rats (Envigo Harlan) were housed individually and were involved in the experiment only during the 12h dark cycle. All animal care and housing procedures complied with National Institutes of Health guidelines and followed protocols approved by the Institutional Animal Care and Use Committee at Johns Hopkins University.

### Dome apparatus

The experiment was conducted in the “Dome,” a planetarium-style virtual reality environment made with fiberglass hemisphere shells (Immersive Display Group, Essex, UK) as described previously^17, 18^. This apparatus consisted of a hemispherical shell (2.3 m inner diameter) onto which visual landmarks were projected by a ceiling-mounted beam projector (Sony VPL-FH30). To provide constant, non-directional illumination, a ring of light was projected near the top of the Dome throughout each session, even when landmarks were absent. Rats were constrained to move in a counter-clockwise direction on an annular table (152.4 cm outer diameter, 45.7 cm inner diameter). This was achieved by attaching the rat’s body harness (Coulbourn Instruments) to a carbon fiber rod that rotated around the center of the table. This radial boom arm also served as a mount for other experimental components, such as infrared lights, feeding tubes, and the recording tether. The tether from the rat’s recording implant (hyperdrive) was connected to a commutator (PSR-36, Neuralynx) mounted at the center of the apparatus, beneath the hemispherical mirror and table. The angular position of the rat was tracked in real-time using two optical encoders: one attached to the commutator drum (Hohner, series INSQ) and another built into the commutator itself. An overhead camera was mounted at the top of the dome to monitor the rat’s behavior throughout the session.

### Electrode implantation and neural recording

6 (2 rats) or 12 (3 rats) independently movable tetrodes were assembled into a 3D-printed hyperdrive implant. Stereotaxic surgery was performed to implant the drive at predetermined skull coordinates using established techniques, under ketamine and isoflurane anesthesia. Further details as well as post-mortem histological figures confirming placement over hippocampal CA1 are available in Reference ^17^. After a minimum of four days of post-operative recovery, during which antibiotics were administered, tetrodes were slowly advanced toward CA1. The final position was confirmed by the presence of high-amplitude sharp-wave ripples in the local field potential (LFP) and isolatable single units and later validated by post-mortem histology. During recording sessions, neural signals were acquired using a Cheetah 5 recording system (Neuralynx). Spike data were filtered between 600–6,000 Hz and digitized at 30 kHz, while LFP data from each tetrode were simultaneously filtered between 1–475 Hz and digitized at the same rate.

### Experimental control

The experiment was managed by a data acquisition system (National Instruments, NI PCIe-6259) and a custom software system built on the Robot Operating System (ROS, Open source Robotics Foundation, distributed under the BSD-3-Clause License) running on a Linux computer. This system received the rat’s angular position from optical encoders and generated the visual scene in the dome using OpenGL ^18^. A second computer running the Windows Operating system was used to acquire neural signals. Liquid rewards were delivered automatically at pseudorandom spatial intervals (typically 40-80°) to encourage continuous running. All experimental data, including rat position, visual stimuli, reward locations, and overhead video, were saved for offline analysis.

### Experimental procedure

Pre– and post-session baseline data were recorded for 20 minutes while the rat rested; comparison of these sessions was used to confirm the stability of single-unit recordings offline. At the start of each session, the rat was attached to the harness at a consistent location relative to the landmarks. The experimenter then exited the dome and monitored the session remotely via an overhead camera, intervening only in cases of equipment malfunction, prolonged inactivity, or an unnatural gait. Session duration varied based on the rat’s running performance and the number of laps planned, and sometimes a second session was conducted if viable. To maintain consistent initial conditions, the rat was typically removed from the apparatus only during epoch 4 (when no landmarks were present), except in sessions where landmarks remained stationary throughout.

The movement of the visual landmarks was dynamically controlled according to the rat’s real-time speed, a preset final landmark gain (G), and the current experimental epoch according to the equation (Landmark speed) = (1 – G) × (Rat speed). Thus, when G > 1, the landmarks were counter-rotated to simulate the animal moving faster than it was, and when 0 < G < 1, the landmarks were rotated in the same direction, albeit at a slower speed than the animal. The landmark behavior was defined as follows:

Epoch 1: Three landmarks were projected and remained stationary (G = 1).

Epoch 2: The landmark gain was linearly ramped from 1 to the final target gain (G_final_). Epoch 3: The landmarks moved at a constant gain G, determined by the rat’s speed.

Epoch 4: The landmarks were not present.

### Estimation of hippocampal gain (*H*)

Standard population decoding techniques were unsuitable for our experiments due to significant remapping events (i.e., place cells appearing and disappearing) that occurred continuously during gain manipulations. To overcome this, we developed a spectral decoding method that is robust to such changes in the active cell population; see our prior reports for details on this gain estimation technique ^17, 19^. This technique leverages the periodic firing of place cells on the circular track to measure the spatial frequency of the population representation, which we term the hippocampal gain, *H*. Conceptually, *H* reflects how many times a place field repeats per physical lap. For a typical place cell with one field, *H* should be 1. If place fields are controlled by landmarks moving at an experimental gain *G*, then *H* should approximate *G*. For example, if landmarks complete one cycle every two laps (*G* = 0.5), we expect *H* to be approximately 0.5. We computed a spatial spectrogram for each unit’s firing rate, analyzing spatial frequencies from 0.16 to 6 cycles per lap using a 12-lap sliding window.

## Data analysis

Hippocampal gain (*H*, derived from hippocampal neuronal activity), landmark gain (*G*, calculated from the real-time position of visual landmarks), and the timestamps for the beginning and end of each epoch were taken directly from the previous study by Jayakumar et al.^17^. This section will therefore focus exclusively on the additional analyses performed for the present study. All analyses were conducted using R (version 4.4.3; R Foundation, Vienna, Austria) and MATLAB (version 2024b; MathWorks Inc., Natick, MA).

### Behavioral analysis

The rat’s position, in degrees, was sampled at 100 Hz via an optical encoder attached to the boom arm. To calculate speed, the raw position data were first smoothed with a 1 Hz low-pass filter. Instantaneous speed was then estimated by taking the difference between the positions at the start and end of a 200 ms sliding window (100 ms before and 100 ms after each sample) and dividing by the window duration, following the method described by Kropff and colleagues^28^. Similarly, instantaneous acceleration was estimated by taking the difference between the velocities at the start and end of a 200 ms sliding window (100 ms before and 100 ms after each sample) and dividing by the window duration.

Both speed and acceleration were calculated at 15 ms sampling intervals (sampling rate ≈ 66.7 Hz). To exclude periods of immobility or slow movement, time points where the rat’s speed was below 10°/s (≈ 11.5 cm/s) were excluded from all analyses, following a previous paper^28^.

### LFP processing and theta metrics

Local field potential (LFP) data, recorded at 30 kHz from each tetrode, were downsampled to 500 Hz and band-pass filtered between 5 and 11 Hz using the bz_Filter function from the BUZCODE toolbox to isolate the theta band^61^. For each tetrode, the amplitude and frequency of the theta-filtered signal were extracted using the Hilbert transform. These values were then smoothed by averaging over a 120 ms sliding window (60 ms before and 60 ms after each time point), following a previous paper^28^. The window was advanced in 15 ms steps, resulting in a 75% overlap between adjacent windows and a continuous estimate of theta dynamics. To obtain a single representative measure for each session, the median of the theta frequency and theta amplitude values was taken across all tetrodes from which hippocampal neurons were recorded. To minimize site-specific differences, theta amplitude was first z-scored within each tetrode across the entire session before the median was calculated. This median-based approach was used to minimize variance arising from differences in recording location. Theta metrics derived from periods when the rat’s speed was below 10°/s were excluded from analysis.

Unless specified otherwise, all analyses of theta dynamics were conducted using a 3-lap sliding window. This window was advanced in 0.1 lap steps. The theta frequency and amplitude within each window were averaged to yield a single representative value for that spatial point. This 3-lap window size was chosen because the minimum epoch 1 size was 3.9 laps; if the window size were bigger than 3 laps, no window would include pure epoch 1 data. This window size differs from the 12-lap window used for hippocampal gain used for the previous study^17^.

Three key reference points were used for analyzing theta dynamics:

1) Baseline: The value measured at a point centered 1.5 laps before the end of epoch 1. This represents the final theta metric before the gain manipulation began.
2) Landmark-off: The value measured at a point centered 1.5 laps before the landmark-off event. This represents the final theta metric before landmarks were removed.
3) Minimum point: The global minimum of theta frequency found between the ‘Baseline’ and ‘Landmark-off’ points. This represents a turning point of theta frequency change in most sessions.

Based on these points, the following metrics were defined:

1) *dTheta* = Landmark-off theta frequency – Baseline theta frequency. This metric quantifies the net change in theta frequency during the period of gain manipulation and stabilization. A dTheta value of 0 indicates that the theta frequency had fully recovered to its baseline level by the time the landmarks were turned off.
2) *theta ratio* =

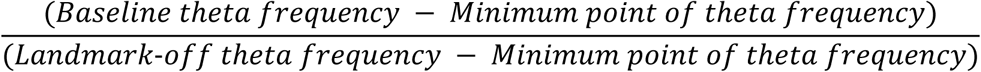 Considering the observed V-shaped modulation of theta frequency, this metric quantifies the degree of frequency recovery relative to its maximal drop. A theta ratio of 1 indicates perfect recovery, whereas a ratio greater than 1 reflects incomplete recovery of theta frequency.
3) *dH*: This metric was defined as the difference between the hippocampal gain measured 6 laps before the landmark-off event (*H*_final_), which is the last landmark-driven gain, and 6 laps after the landmark-off event (*H*_recal_), which is the first path-integration-driven gain. Given that hippocampal gain tended to revert to the gain of 1 after the landmark-off event, *dH* was calculated as

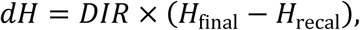

where *DIR* is a directional indicator, defined as +1 for gain-up sessions and −1 for gain-down sessions. In both gain-up and gain-down sessions, a *dH* of 0 represents perfect recalibration, while a positive value indicates incomplete recalibration.

### Session selection and definition

From the 72 pre-screened sessions reported in the previous study^17^, sessions were included for analysis if they met the following criteria:

1) The experimental gain was modulated in only one direction (i.e., gain-up or gain-down), and the final experimental gain was within the range of [0.1, 1.8].
2) The presence of action potentials and the presence of stable spatial tuning, as determined by qualitative inspection, were required in the 12 laps preceding and following the landmark-off event. Sessions were also required to have at least 12 laps of data recorded after the landmark-off event.
3) The mean gain ratio, defined as the hippocampal gain divided by the landmark gain in the period before the landmark-off event, was less than 1.1.

Sessions that met all of the above criteria were classified as ‘landmark-controlled’ (n = 21 gain-up [rat 515: 3, rat 576: 3, rat 637: 3, rat 638: 6, rat 692: 6], 14 gain-down [rat 515: 3, rat 576: 4, rat 637: 2, rat 638: 1, rat 692: 4]). Sessions that met the first two criteria but failed the third were classified as ‘landmark-failure’ sessions and were analyzed separately (n = 7 [rat 637: 3, rat 638: 1, rat 692: 3]). For these landmark-failure sessions, the ‘landmark-failure point’ was operationally defined as the last point at which the difference between the hippocampal gain and the landmark gain was less than 0.015 before their subsequent divergence. Unless specified otherwise, all reported analyses were performed on the landmark-controlled sessions.

### Overlay plot construction and statistical metrics

Overlay plots (Fig. 4c, S1a, S1d, S2c, and S4c, S6a) were constructed by aligning the theta frequency trace from each session to either the end of epoch 2 or the landmark-failure point. To generate a mean frequency trace that was aligned relative to a given frame, data points were averaged across all sessions, gain-up sessions, or gain-down sessions, with the condition that each lap point must include data from at least three sessions. In this analysis, sessions were averaged regardless of rat identity; however, session traces for each individual rat were reported separately (Fig. S1d).

Because the number of trials per epoch varied across sessions, applying a single continuous scaling to all trials in a session would cause misalignment of epoch boundaries. To address this issue, we performed scaling separately for each epoch. For the rescaled overlay plots (Fig. 1c, 4b, and S2b), the theta frequency traces were normalized along the lap axis such that the boundaries of each epoch were aligned across sessions. To achieve this, epochs 2 through 4 were each uniformly sampled into nine equal segments, while epoch 1 was sampled into two segments due to its shorter average duration. This visualization method allows for an intuitive comparison of theta frequency dynamics across epochs, regardless of variations in session or epoch length.

In the case of landmark-failure sessions, the landmark failure event was added as a reference point as well as epoch boundaries (Fig. 4b). Therefore, epoch 2 was segmented into 8 equal segments instead of 9 to observe the frequency modulation by the landmark failure event (LF). In this plot, the first four segments represent the range between the epoch 2 start and landmark-failure event, while the last four segments represent the range between the landmark-failure event and the end of epoch 2.

To quantitatively assess the observed V-shaped modulation of the average theta frequency trace, we used the following two metrics for overlay plots:

*Slope.* We employed continuous piecewise regression to fit two linear segments that best described the average theta frequency trace. The slopes of two regression lines were estimated using the lsqcurvefit function (Optimization Toolbox, MATLAB) with the trust-region-reflective algorithm. The model was defined with five parameters:

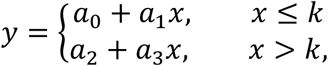

where *y* represents theta frequency and *x* represents laps. The location of the break point *k*, the intercept of the first line *a*_0_, the slope of the first line *a*_1_, and the slope of the second line *a*_3_, were used as free parameters. The intercept of the second line (*a*_2_) was determined by the constraint of continuity and thus was not a free parameter. Initial guesses for *a*_0_, *a*_1_, *a*_3_, and *k* were 0, –1, 1, and the center of the data range, respectively. During optimization, the breakpoint was constrained by upper and lower bounds to remain within the data range. This breakpoint was used solely for calculating the slopes, as the global minimum point in the data was used to quantify the minimum value of the V-shape. The slope difference (Δ*m*), defined as a_3_ – a_1_, quantifies the difference between the two slopes and serves as an indicator of how closely the pattern approximates an ideal V-shape.

*Distance*. This was defined as the lap-wise distance between the global minimum (global maximum for theta amplitude analysis) point in the average theta frequency trace and the reference point (end of epoch 2 or landmark failure event) to quantify their relative positions.

For the model comparison, we employed the Bayesian Information Criterion (BIC), defined as follows:

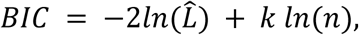

where *k* represents the number of parameters, 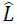 denotes the maximum likelihood, and *n* indicates the sample size. To evaluate whether the fit of the V-shaped profile observed in the average theta frequency trace was superior to a standard linear model, we adopted the Δ*BIC* metric, defined as the *BIC* of the piecewise model minus the *BIC* of the linear model. We utilized this metric for two primary reasons. First, break points in some shuffled data were located at the extreme edge of the data range, yielding high slope differences despite being indistinguishable from linear fits. This metric allows more rigorous identification of an ideal V-shape while accounting for these edge-case artifacts. Second, when comparing empirical data to shuffled distributions (Fig. S1c) or evaluating different alignment references for the same condition (Fig. 4c and S6a), the number of samples used to construct each piecewise model often varies. We minimized the influence of the sample size by assessing the relative improvement of the piecewise fit over the linear model within each dataset.

### Shuffling procedure

To assess the statistical significance of our metrics, we performed a circular shuffling procedure. For each of the 10,000 iterations, the theta frequency trace from each session was circularly shifted by a random amount and a new average trace was computed. This circular shifting was performed only in the range of data in which each data point contains at least 3 samples. The shift range was set from 0 to 1 full cycle to ensure maximal randomization. This process generated null distributions for the slope change and the distance of the minimum point from the reference point. P-values were calculated as the proportion of samples from the null distribution that were more extreme than the observed value (i.e., above the 95^th^ percentile or below the 5^th^ percentile). All methods described above were applied identically to the analysis of theta amplitude.

### Linear mixed-effects modeling

To explain *dH* at the population level, linear mixed-effects models were constructed using R (version 4.4.3; R Foundation, Vienna, Austria), RStudio (Posit Software, Boston, MA), the lme4 package ^26^, and lmerTest package ^62^ to account for the repeated measures within subject. Unless otherwise specified, the general structure of the linear models was as follows:

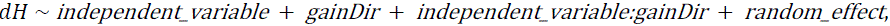

where *gainDir* is a dummy-coded variable (0 or 1) representing the session’s gain direction (gain up or gain down). The *independent_variable* was specified as *dTheta*, *theta ratio*, locomotion-adjusted *dTheta*, or locomotion-adjusted *theta ratio*, as appropriate for each model.

For every analysis, we tested three different random effect structures:

Full random-effect model:(1 + *independent*_*variable*|*rat*)

Random-slope-only model: (0 + *independent*_*variable*|*rat*)

Random-intercept-only model: (1|*rat*)

where *rat* refers to the animal’s identity. We report the coefficient for the *independent*_*variable* (estimate of coefficient and its 95% confidence interval). Normality was assessed using the Shapiro-Wilk test. Since the residual of a model did not meet the assumption of normality in some models, 95% confidence intervals were estimated using a non-parametric bootstrapping procedure (1,000 resamples) to ensure robust statistical inference. Bootstrapped confidence intervals were obtained using the confint.merMod function from the lme4 package for linear mixed-effects models and the boot function from the car package. Coefficients were considered statistically significant when the 95% confidence interval did not include zero.

The results from the random-intercept-only model were reported in the main text because this model yielded the lowest BIC value in most analyses. The results from the other random-effect models were documented in the supplementary figures for reference. However, when the random-intercept-only model resulted in a singular fit, another random-effect model that showed the lowest BIC was presented in the main text and the random-intercept-model was reported in the supplementary figure. Most full random-effect models resulted in a singular fit (or nearly singular fit) likely due to an insufficient number of sessions and high covariance between random effects; nevertheless, the statistical outputs are reported along with a note of this issue. Random-slope-only models were also reported for reference purpose.

For landmark-failure sessions, because all landmark-failure sessions were gain-down sessions, the *gainDir* fixed effect and its interaction term were removed from the model.

Otherwise, the analysis proceeded in the same manner as for the landmark-controlled sessions.

### Analysis of locomotion-adjusted theta frequency

To investigate whether the relationship between theta dynamics (*dTheta*, *theta ratio*) and gain mismatch was merely an artifact of locomotion factors (speed and acceleration), we performed a decomposition analysis. We defined a set of 3-lap sliding windows (1.5 laps before and after each sampling point) with a 2.9-lap overlap between adjacent windows, covering epochs 1 through 3. These windows matched those used to calculate the theta samples for the *dTheta* and *theta ratio* analyses. Within each window, we constructed the following linear model using the theta frequency, speed, and acceleration data:

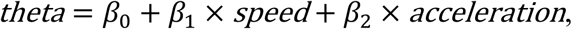

where *theta* is the z-transformed theta frequency, and β_1_ and β_2_ are the coefficients for speed and acceleration, respectively. The intercept β_0_, which represents the portion of theta frequency not explained by these locomotion factors, was termed the ‘locomotion-adjusted theta frequency’. All subsequent analyses addressed above for the theta frequency were executed with identical procedures, with the exception that the theta frequency, speed, and acceleration were sampled at 33.33Hz. To visualize the relative contribution of each locomotion factor to the theta frequency within each sliding window, we plotted the window-averaged speed multiplied by its corresponding coefficient, the window-averaged acceleration multiplied by its corresponding coefficient, and the intercept for a representative example session (Fig. S4b).

### Individual session analysis

To determine whether theta frequency could predict the lap-by-lap error between path integration gain and landmark-controlled hippocampal gain within individual sessions, we performed the following multi-step analysis:

1. To estimate the lap-by-lap path-integration gain from the hippocampal gain, which is not directly measurable while landmarks are present, we made the following assumptions:
  a. The path-integration gain during epoch 1, when landmarks were stationary, was assumed to be the baseline gain of 1. Therefore, the path-integration gain at the end of epoch 1 was set to 1.
  b. We assumed that the path-integration gain at the 6^th^ lap before the landmark-off event (end of epoch 3) was identical to the first measurable path-integration gain, which was represented by the hippocampal gain at the 6^th^ lap after the landmark-off event (beginning of epoch 4). This assumes that the path-integration gain was not changed by the landmark-off event.
  c. Based on the assumption that path-integration gain changed continuously, we estimated the lap-by-lap values via linear interpolation.
2. Calculation of lap-by-lap gain error. The lap-by-lap gain error was calculated as the difference between the estimated path-integration gain (*P*) and the hippocampal gain (*H*) as below:

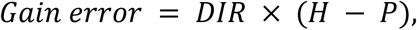

where *DIR* is a directional indicator, defined as +1 for gain-up sessions and −1 for gain-down sessions. To visually demonstrate the inverse relationship between this gain error and theta frequency, representative examples are shown in Figure 3b, where the gain error (blue) is overlaid with the inverted theta frequency (purple).
3. Linear modeling of gain error. For each session, we then tested whether theta frequency could significantly predict the gain error on a lap-by-lap basis by fitting the following linear model:

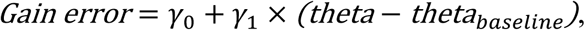

where (theta − theta_*baseline*_) is the theta frequency minus the baseline theta frequency. Because this model was fitted to lap-by-lap data within a single session, animal identity was not included as a random effect. The analysis was restricted to data prior to the 6th lap before the landmark-off event, where the final landmark-driven hippocampal gain was measured. These linear models were fitted independently for each session; therefore, rat identity for each rat was not considered. We calculated and reported the 95% confidence interval for the slope coefficient (*y*_1_) for all sessions with the bootstrapping procedure described above. To determine if the proportion of sessions with a significant relationship (p < 0.05) exceeded what would be expected by chance, we performed a binomial test. The chance level for a significant p value was set to 5% when testing whether the number of significant sessions exceeded chance, and to 50% when testing whether the number of negative sessions exceeded the number of positive sessions.

## Continuous attractor neural network model

The model used in this study was adapted from a previously published model by Chu and colleagues^30^. A detailed description of the model architecture and its parameters can be found in Chu et al.^30^ and in Supplementary Table 1.

Our research group previously used this model to gain insight into theta phase precession and phase procession at the single-cell level during gain recalibration, so some parameters were adopted from this paper^31^. Our current efforts focused on the model’s capacity to describe how the ensemble representation of the current position in the hippocampus, the ‘bump’ of activity, oscillates at a theta frequency. In this model, the bump oscillation is generated by population activity. Accordingly, we utilized this model to observe how the oscillation frequency of this bump changes when different landmark gains and path integration gains, analogous to those in an actual experiment, are given in the simulation. The equations we directly adopted from Chu et al.^30^ models are described below. The synaptic drive onto the ring attractor was calculated as

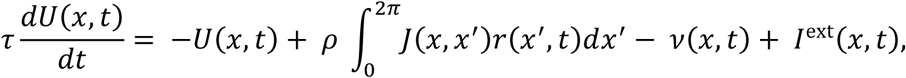

where U(x, t) represents synaptic inputs to the place cell at location x and time t, τ denotes the time constant of the synaptic dynamics, ρ represents neuron density, *J*(*x*, *x*’) represents connection weight from a neuron *x*′ to a neuron at *x*, *r* represents firing rate, *v*(*x*, *t*) represents an adaptation effect, which destabilizes the attractor state, and *I*^ext^(*x*, *t*) represents external input. The firing rate function is defined as

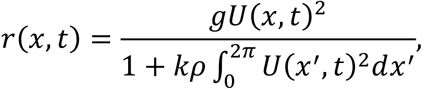

where g denotes a gain factor and k denotes the strength of the global inhibition. Connection weights are defined as

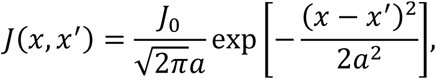

where J_0_ represents the strength of the recurrent connections, and *a* denotes the range of neuronal interaction. The adaptation dynamics are defined as

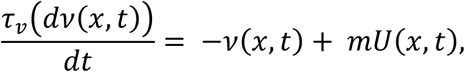

where τ_v_ denotes the time constant of *v*, and m denotes the adaptation strength. The external input is defined as

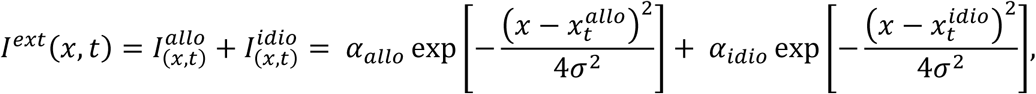

where α_*allo*_ and α_*idio*_ denote the strengths of the allocentric input, 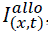, and the idiothetic input, 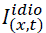, respectively. The parameter σ parameterizes the width of the external input. The terms 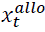 and 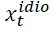 denote the rat’s actual location and its estimated location from path integration, respectively, at time *t*, in the landmark frame. The rat’s actual location is defined as 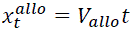, where *V*_*allo*_ denotes the animal’s running speed in the landmark frame. By nature, the estimated location by path integration accumulates error, and this error is corrected by external landmarks ^15^. Thus, the rat’s estimated location by path integration is defined as

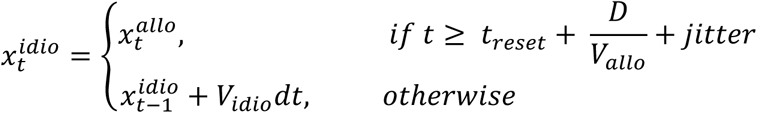

Where *t*_*reset*_ denotes the last correction time, *D* is a distance threshold in the allocentric frame at which accumulated path-integration error is corrected, and *V*_*idio*_ denotes the speed estimated by path integration in landmark frame. A random jitter, ranging from –100 ms to 100 ms, was introduced to the reset interval to prevent the error correction timing from becoming phase-locked to a specific phase of the oscillation. Let *V* denote the velocity of the animal in the laboratory frame, in which case *V*_*idio*_ = *P V* and *V*_*allo*_ = *G V* and thus, we can eliminate *V*:

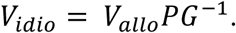

To demonstrate the general trend of theta frequency by input amplitude, we fixed both *G* and *P* to 1 so as not to allow discrepancy of gains and measured the oscillation frequency in both high input amplitude (*alpha* = 1.1) condition and low input amplitude (*alpha* = 0.08) condition (Figure 6d).

To generate results comparable to our experimental conditions, we constrained the range of *G* to [0.3, 1.7] and *P* to [1, *G*] (or [*G*, 1] if *G* < 1). We then ran simulations to determine the oscillation of the ‘bump’, representing the current position encoded by the hippocampal population. Each simulation was run for 60 seconds. The initial 6 seconds (transient behavior) were discarded, and the subsequent 54 seconds of the bump’s position were tracked for analysis. The frequency of the bump oscillation was calculated using a Discrete Fourier Transform using Matlab’s fft command. To quantify the hippocampal gain (*H*), we performed a linear fit on the bump’s trajectory over time; the resulting bump velocity was then multiplied by 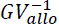.

We calculated the change in theta frequency (*dTheta*) by subtracting the baseline theta frequency (obtained under the *G* = 1, *P* = 1 condition, which mimics epoch 1) from the frequency measured in each *G* and *P* condition. The gain difference between landmark-controlled hippocampus gain and path-integration gain (*dG*) was defined as

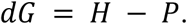

*dG* was then assessed using the fitlm function in MATLAB with the following linear model:

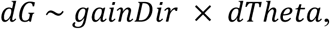

where *gainDir* is a directional indicator, defined as +1 for gain-up sessions and −1 for gain-down sessions.

To generate the representative theta frequency traces shown in Figure 6f, we first established the final hippocampal and path-integration gains and constructed their temporal profiles. Similar to the experimental protocol, the simulation was divided into three epochs with durations of 60, 180, and 360 seconds, respectively. Hippocampal gain and oscillation frequency were calculated as described above; however, instead of using the entire range, we employed a 12-lap sliding window. This window moved in increments of 0.1 laps, and the hippocampal gain and oscillation frequency were computed for each window and assigned to its central lap for plotting. Consistent with previous modeling work, the MATLAB fft function was used to estimate the oscillation frequency ^30^. To enhance the detection of subtle frequency modulations, a Hanning window was applied to the detrended bump oscillation in each window, followed by zero-padding to 16 times the original window length to increase spectral interpolation.

## Declaration of Generative AI and AI-assisted technologies

During the preparation of this manuscript, the authors utilized Gemini 3 and 3.1 (Google) to enhance the technical quality and clarity of the presentation. AI assistance was strictly limited to the following areas: (1) language editing and refinement of the text and (2) implementation of author-defined algorithms into scripts, including improvements to code readability and data presentation. After using this tool/service, the authors reviewed and edited the content as needed and they take full responsibility for the content of the publication.

## Acknowledgements

We thank Gorkem Secer, Yotaro Sueoka, Zilong Ji, Tianhao Chu, and the members of Knierim and Cowan laboratories for helpful discussions. This research was supported in part by National Institutes of Health grants R01 MH079511 (H.T.B., J.J.K.), R01 NS102537 (J.J.K., N.J.C. and F.S.), R01 MH118926 (J.J.K. and N.J.C.), R21 NS095075 (J.J.K. and N.J.C.), and R01 NS039456 (J.J.K.); the Johns Hopkins Kavli Neuroscience Discovery Institute (M.S.M.); and a Johns Hopkins University Discovery Award (N.J.C., J.J.K.).

