## Supplementary data for "Hippocampal theta frequency as a readout of path-integration recalibration"

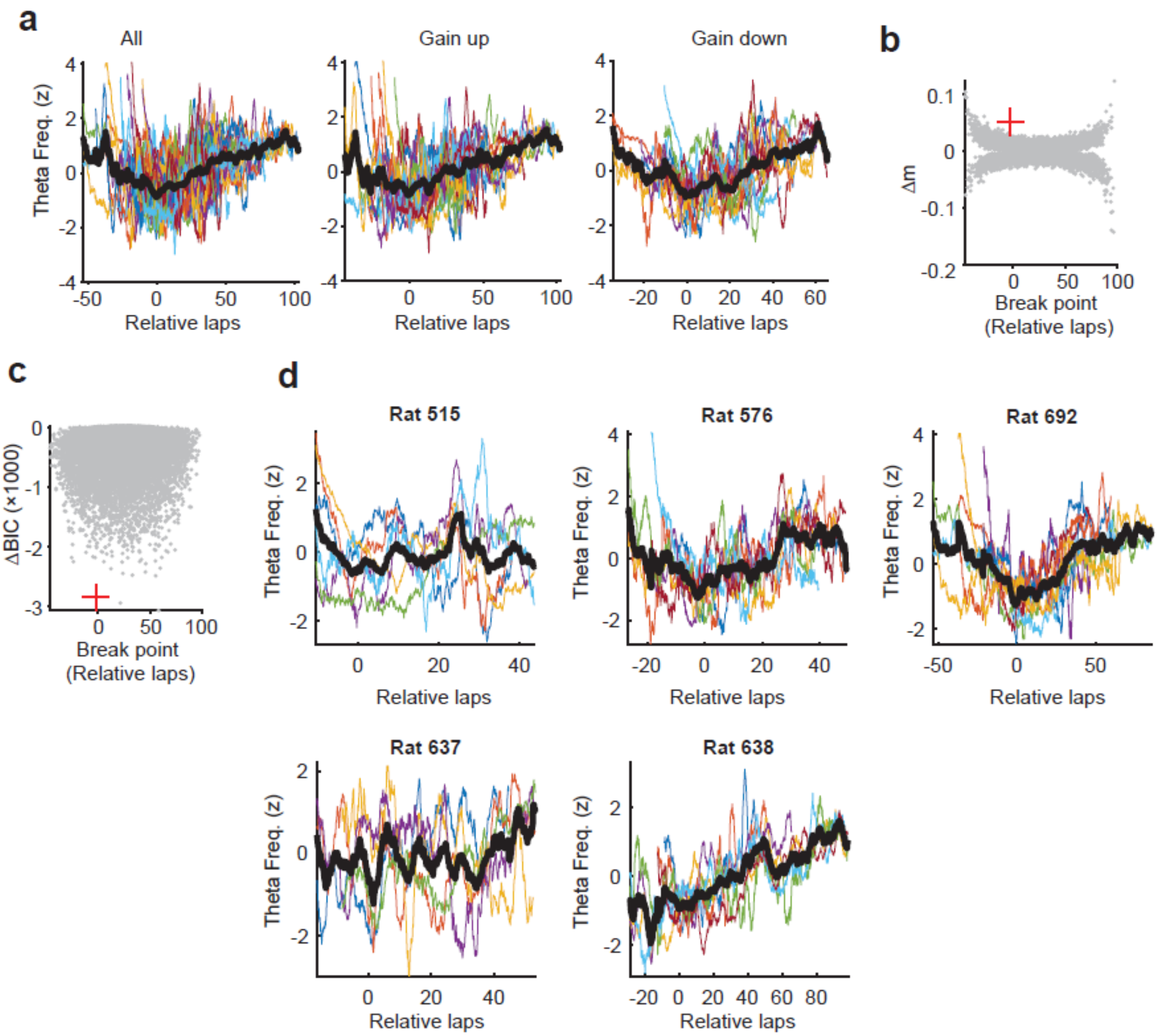


Supplementary Figure 1. Non-rescaled theta frequency dynamics and statistical distribution.

a. Overlay of theta frequency traces aligned to the end of epoch 2. Theta frequency traces from individual sessions are displayed without temporal rescaling. The x-axis indicates relative laps from the end of epoch 2. Plots are shown for all sessions (left), Gain up sessions (middle), and Gain down sessions (right). Individual traces are color-coded, with the group average in bold black. The average was computed only for spatial bins containing data from at least three sessions.

b. Distribution of breakpoints (i.e., the point where the two lines of the piecewise linear regression meet) and slope differences (Δm) for shuffled data. The zero point on the x-axis denotes the end of epoch 2. A higher Δm indicates a sharper V-shaped profile. The red cross marks the breakpoint and Δm derived from the measured data; this point lies outside the distribution of the shuffled data.

c. Distribution of breakpoints and ΔBIC for shuffled data. ΔBIC is defined as the difference between the BIC of the piecewise regression model and that of a simple linear model. A more negative ΔBIC indicates that the piecewise fit explains the data better than the linear fit. The red cross marks the experimental data point, which lies outside the distribution of the shuffled data. Note that while panel b shows some shuffled samples with breakpoints located near the ends of the x axis exhibit a sharper Δm than the measured data, these samples show a ΔBIC closer to zero in this panel. This suggests that they do not deviate substantially from the linear model compared to the observed data.

d. Individual animal theta frequency dynamics. For each rat, colored lines represent individual sessions, and the bold black line indicates the session average. Averaging was restricted to laps with at least three sessions to prevent individual sessions from disproportionately influencing the mean. The x-axis represents relative laps from the end of epoch 2. Although the data are noisier within each rat, the global minimum point is located near zero in rats 515, 576, and 692. While this pattern is less pronounced in rats 637 and 638, a local minimum near the end of epoch 2 occurs in these rats as well.

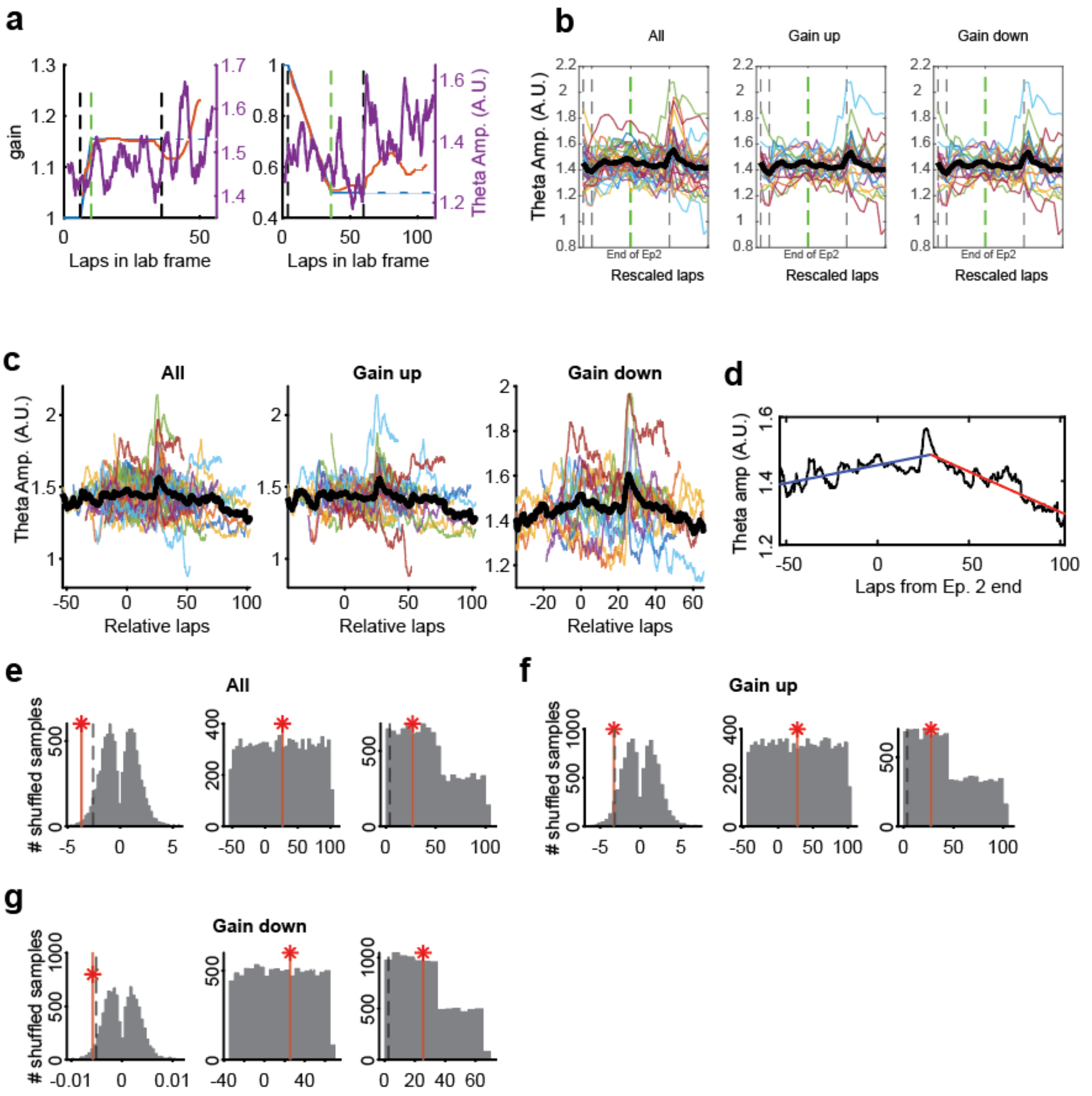


Supplementary Figure 2. Theta amplitude dynamics during the experiment.

a. Representative examples of theta amplitude dynamics from two single sessions. These examples are presented analogous to the frequency examples in Fig. 1b. Blue, orange, and purple lines indicate experiment gain, hippocampal gain, and theta amplitude, respectively. As theta amplitude is known to vary depending on the recording site [1], we first z-scored the filtered theta band signal from each tetrode to minimize site-specific differences before computing the median amplitude for the session. Each vertical dashed line shows the edge of each epoch.

b. Overlay of theta amplitude traces with each epoch rescaled to a uniform duration, analogous to the frequency analysis in Fig. 1c. Theta amplitude showed no discernible change over the course of the experiment, except for a sharp, transient increase at the landmark-off event.

c. Overlay of theta amplitude traces aligned to the end of epoch 2, without temporal rescaling. This visualization is analogous to the theta frequency plot in Supplementary Fig. 1a.

d. Quantification of the V-shaped profile of the theta amplitude using piecewise regression on the average trace, analogous to Fig. 1d.

e. Statistical validation of theta amplitude V-shape metrics via shuffling tests. The observed value from the group average is marked with a red star and a red vertical line, while the gray dashed lines represents the 5$^{th}$ percentile. The histograms show the distributions for shuffled data representing $\Delta m$ (left, -0.0037, p = 0.0125), the global maximum location (center), and its distance from the end of epoch 2 (right, 26.8 laps, p = 0.3450). Because the upper angle is >180°, the peak of the averaged theta amplitude trace, rather than the valley, was measured as the reference point. Even though the slope change is significantly different from the shuffled data, it appears to be associated with the abrupt surge at the landmark-off event rather than a sustained modulation (see panel b).

f-g Distributions of shuffled data for Gain up (f) and Gain down (g) sessions. Plots show $\Delta m$ (left), peak location (center) and peak distance (right), analogous to (e). For Gain up sessions, $\Delta m$ = -0.0033 (p=0.0458) and distance = 27.5 laps (p=0.3740). For Gain down sessions, $\Delta m$ = -0.0057 (p=0.0329) and distance = 25.5 laps (p=0.5120).

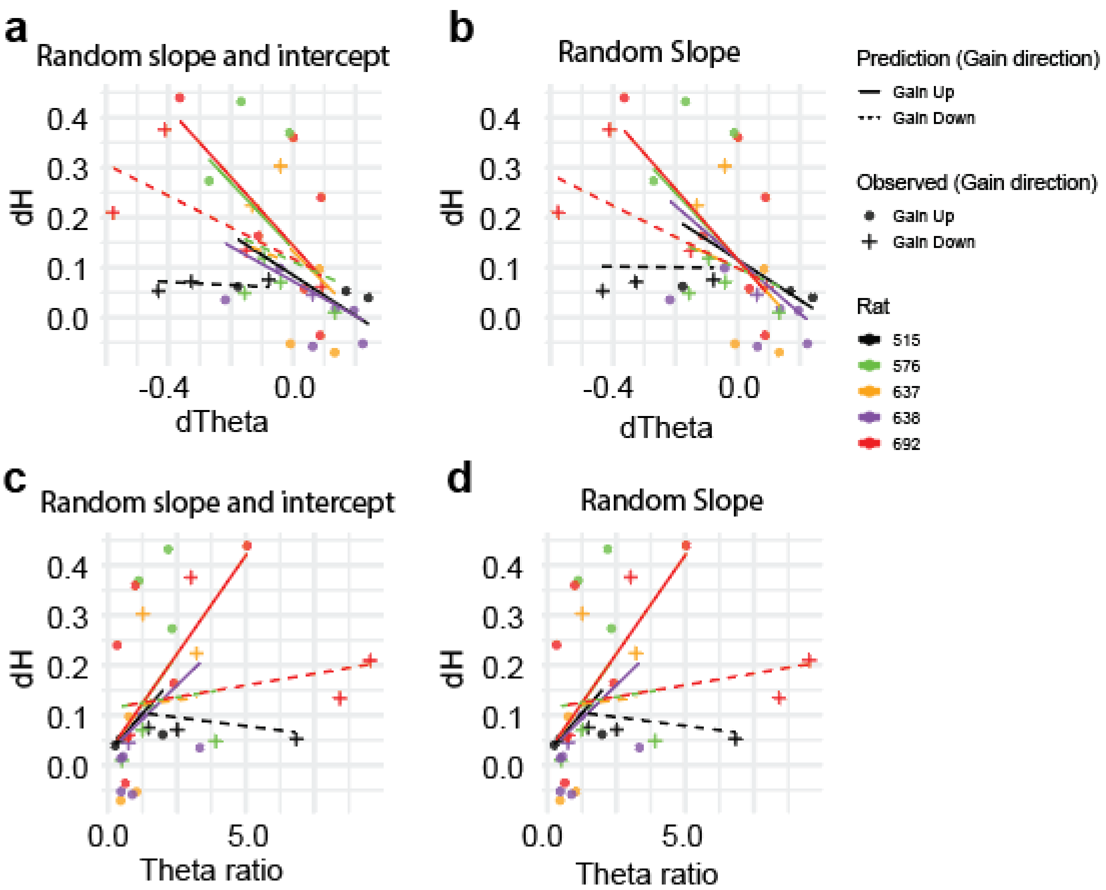


Supplementary Figure 3. Theta frequency dynamics predict the conflict between hippocampal gain and path-integration gain. In all mixed-effects models, gain direction was treated as a fixed effect and rat identity was used as a random effect unless specified otherwise.

a-b. dTheta as a predictor of dH

a. Full model with random intercepts and slopes for dTheta. This model resulted in a singular fit, indicated by a perfect (-1.00) or near-perfect correlation between the random effects. A singular fit suggests that the structure of random effects is too complex for the available data, making it difficult to reliably estimate the random effects, often due to an insufficient number of samples or an extreme correlation between the random intercept and slope [2]. Therefore, this model was considered unreliable for interpretation ($\beta$ = -0.547, CI = [-0.898,-0.187], $p < 0.05$, BIC = -10.6).

b. Random-slope-only model for dTheta. While this model had a slightly lower BIC than the random-intercept-only model presented in the main text ($\Delta$BIC = 0.2), the difference was negligible and did not alter the interpretation. For consistency with other analyses, in which random-intercept-only models have the lowest BIC, it is reported here ($\beta$ = -0.602, CI = [-0.979,-0.224], $p < 0.05$, BIC = -15.8).

c-d. The theta ratio as a predictor of dH

c. Full model with random intercepts and slopes for the theta ratio. Similar to (a), this model resulted in a singular fit and was not used for interpretation. A significant main effect ($\beta = 0.065$, CI = [0.018,0.111], $p < 0.05$, BIC = 3.6) and a significant interaction with gain direction ($\beta = -0.063$, CI = [-1.171,-0.010], $p < 0.05$) were observed.
d. Random-slope-only model for the theta ratio. This model also showed a significant main effect of the predictor ($\beta = 0.070$, CI = [0.016,0.123], $p < 0.05$, BIC = -2.2) and a significant interaction with gain direction ($\beta = -0.070$, CI = [-0.128,-0.016], $p < 0.05$).

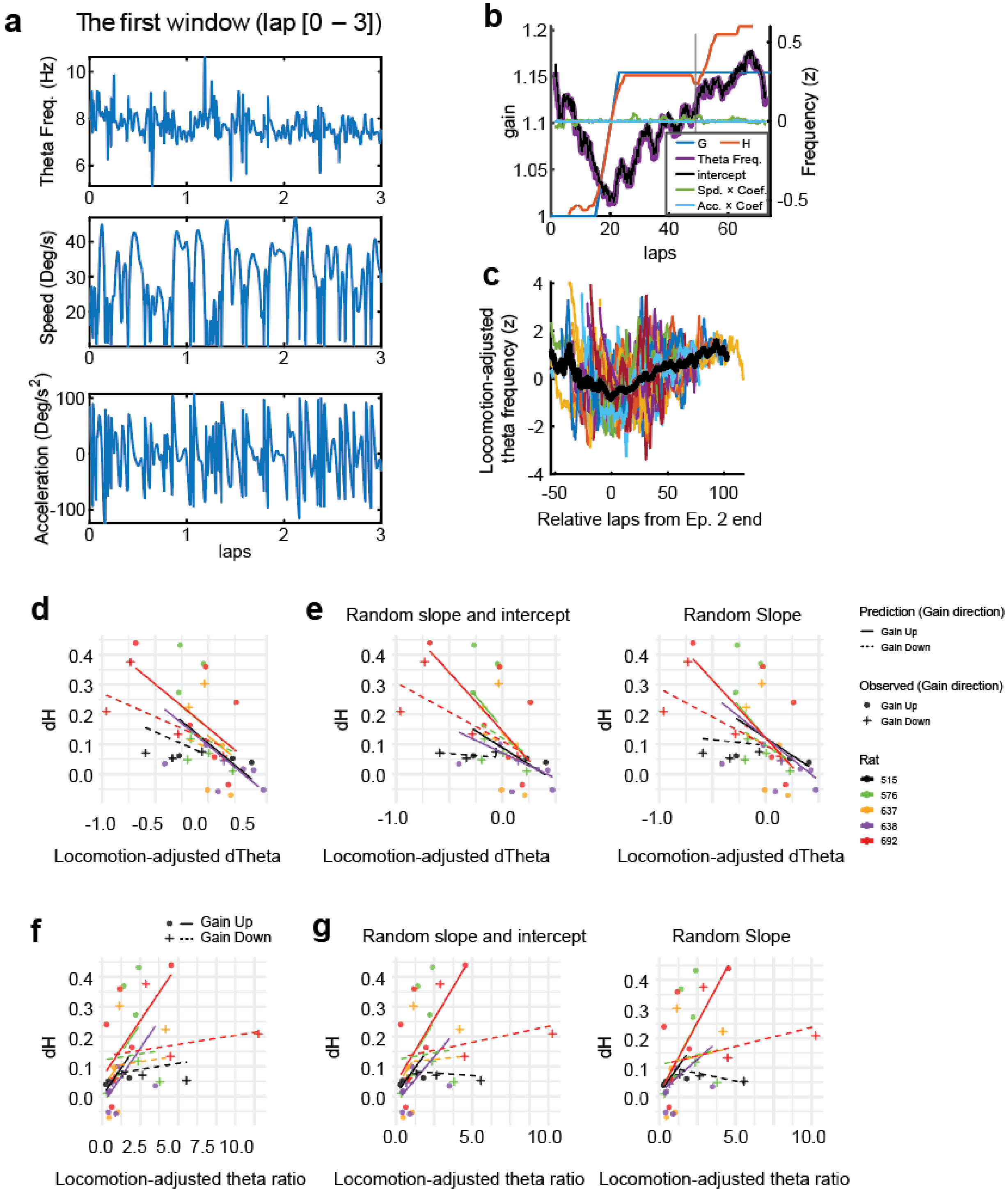


Supplementary Figure 4. The ability of theta frequency to predict the magnitude of gain recalibration is not confounded by effects of speed or acceleration

Hippocampal theta frequency is known to be modulated by locomotor parameters such as velocity and acceleration, as well as by contextual factors such as novelty and mismatch[3-7]. We thus considered the possibility that changes in theta frequency primarily reflected biased changes in velocity or acceleration during the session rather than mismatches between path-integration gain and experimental gain. Because H is a variable that is measured over a time course of 12 laps, whereas speed and velocity are variables that can vary widely over timescales of seconds, this time-scale inconsistency prevented the use of a regression model in which H could be considered as an independent variable along with speed and acceleration. We thus reasoned that if we fit a linear model to account for the effects of speed and acceleration, the intercept term of that model would reflect the influence of H on theta frequency without the contributions of speed and acceleration. We defined spatial analysis windows encompassing three laps each, and within each window, we fit the following linear model:

$$\text{Theta frequency} = \beta_0 + \beta_1 \times \text{velocity} + \beta_2 \times \text{acceleration},$$

where the intercept term $\beta_0$ represents the locomotion-adjusted theta frequency that is the component not explained by velocity or acceleration, and $\beta_1$ and $\beta 2$ represent the coefficients of velocity and acceleration (Panel a-b). This windowed approach allowed us to preserve fine-grained fluctuations in behavior while quantifying the non-locomotor contributions to theta frequency. Each window contained 3 laps and was placed 0.1 lap apart from neighboring windows, allowing 5.9 laps overlap between neighbors. In each window, linear models were constructed independently, enabling us to estimate locomotion-adjusted theta frequency continuously along the track (Panel a-b). As shown in b, the locomotion-adjusted theta frequency ($\beta_0$) closely paralleled the raw theta frequency. Overlay plots of the locomotion-adjusted theta frequency exhibited a prominent V-shaped pattern, mirroring that of the raw frequency, suggesting that this non-behavioral component preserves key features of the original signal (Panel c).

A mixed-effects model showed that greater decreases in locomotion-adjusted theta frequency predicted larger mismatches between hippocampal gain and path-integration gain ($\beta = -0.294$, CI = [-0.477,-0.116], $p < 0.05$, BIC = -13.8; Panel d and e). Similarly, a higher locomotion-adjusted theta ratio was associated with a higher dH, suggesting that lower recovery of locomotion-adjusted theta frequency is associated with larger gain difference between hippocampus gain and path-integration gain. ($\beta = 0.076$, CI = [0.027,0.128], $p < 0.05$, BIC = -13.8; Panel f and g). These results demonstrate that the ability of theta frequency changes to predict the mismatch between hippocampal gain and path-integration gain is not an artifact of speed or acceleration confounds.

a. Examples of the raw input data (theta frequency, velocity, and acceleration) from the first 3-lap window that was used to compute a single $\beta_0$ value.
b. Decomposition of theta frequency in a representative session. The raw theta frequency (purple) is decomposed into its three main components: the intercept ($\beta_0$, locomotion-adjusted theta frequency; black), the velocity component (velocity $\times$ $\beta_1$; green), and the acceleration component (acceleration $\times$ $\beta_2$; cyan). The locomotion-adjusted component (black) closely tracks the raw theta frequency, while the contributions from locomotion factors are minimal, demonstrating that the V-shaped modulation is not an artifact of running speed or acceleration.
c. Overlay of locomotion-adjusted theta frequency ($\beta_0$) traces from all sessions. Traces are

aligned to the end of epoch 2 (0 on x axis), revealing a consistent V-shaped modulation similar to that of the raw theta frequency. This plot is analogous to the plot in Fig. S1a.

d. The change in locomotion-adjusted theta frequency predicts gain mismatch. This plot is analogous to the plot in Fig. 2c.

e. Locomotion-adjusted dTheta as a predictor of dH. (Left) The full model, including random intercepts and slopes (1 + locomotion-adjusted dTheta | rat), resulted in a singular fit and was considered unreliable ($\beta = -0.318$, CI = [-0.536,-0.108], $p < 0.05$, BIC = -9.0). (Right) The random-slope-only model (0 + locomotion-adjusted dTheta | rat) shows a significant negative relationship, indicating that a greater decrease in theta frequency predicts a larger gain error even after accounting for locomotion ($\beta = -0.346$, CI = [-0.579,-0.122], $p < 0.05$, BIC = -13.7).

f. The theta ratio of locomotion-adjusted theta frequency predicts gain mismatch. This plot is analogous to the plot in Fig. 2d.

g. The locomotion-adjusted theta ratio as a predictor of dH. (Left) The full model resulted in a singular fit and was not used for interpretation. It showed a significant main effect ($\beta = 0.076$, CI = [0.025,0.126], $p < 0.05$, BIC= -9.0). (Right) The random-slope-only model also showed a significant main effect ($\beta = 0.082$, CI = [0.028,0.139], $p < 0.05$, BIC= -13.7).

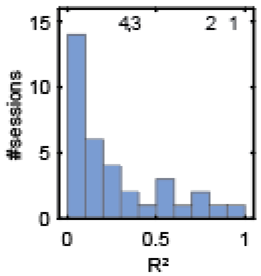


Supplementary Figure 5. Supplementary figure for the individual session analysis.

Distribution of $R^2$ values of Gain error and inverted theta frequency across all landmark controlled sessions (See Figure 3b). The histogram illustrates the $R^2$ distribution for all landmark-controlled sessions, including the representative examples shown in Figure 3a-d (indicated by numbered labels). The number location indicates the corresponding $R^2$ of the example number. This distribution confirms that the selected examples are representative of the entire dataset rather than outliers ($R^2$ of the example sessions: 0.90, 0.78, 0.36, 0.29, respectively).

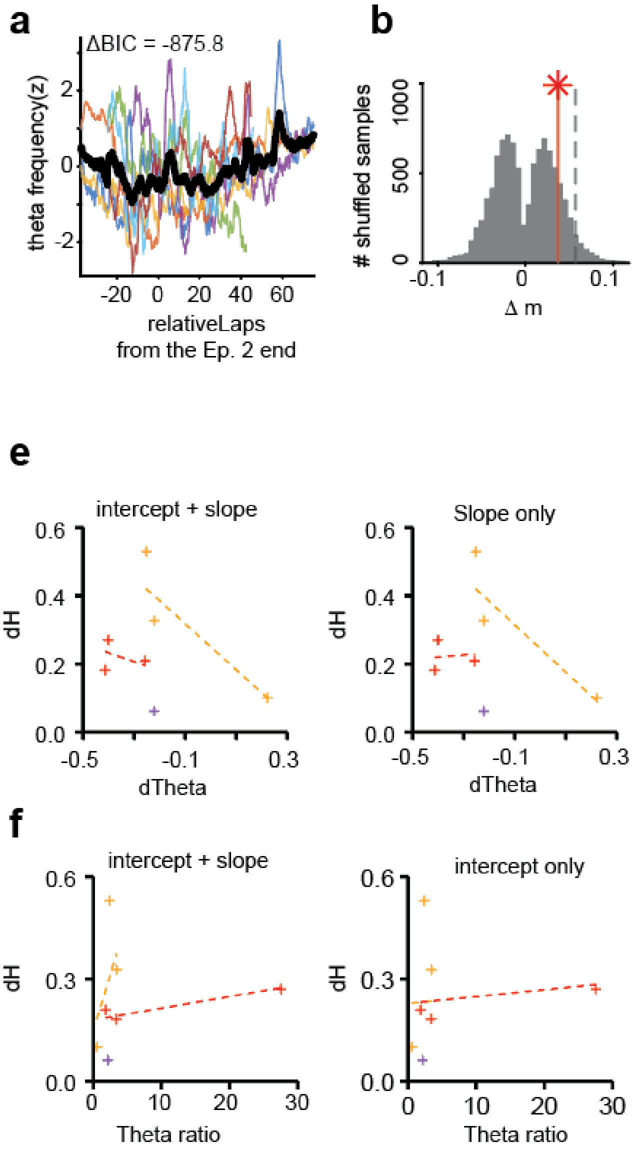


Supplementary Figure 6. Supplementary data of landmark failure sessions.

a. Overlay of theta frequency traces aligned by the end of epoch 2. This plot is analogous to Fig. 1c. ΔBIC, which is defined as (BIC of piecewise fit – BIC of linear fit), is shown in the figure. The large negative ΔBIC value indicates that the piecewise regression model provides a significantly better fit than a simple linear model, supporting the existence of a non-linear frequency shift. and the distance between the lap with the global minimum of theta frequency and the end of epoch 2 was not significantly shorter than chance (12.7 laps, $p = 0.2246$, shuffle test, Panel c and d), confirming that the dynamics in these sessions differed from those in landmark-controlled sessions.

b. Shuffle test for the Δm of the V-shaped theta frequency profile. In contrast to the landmark control sessions, the resulting slope difference (Δm) of the V-shape was not significantly more acute than chance (Δm = 0.037, p = 0.1574, shuffle test). The observed value from the group average is marked with a red star and a vertical line, with the histogram representing the distribution of the shuffled data. The gray dashed line represents $95^{th}$ percentile of shuffled data.

c. Distribution of the lap with the theta frequency global minimum from shuffled data. The x-axis indicates relative laps from the end of epoch 2; the observed value from the group average is marked with a red star and a vertical line.

d. Shuffle test for the distance of the lap with the theta frequency global minimum from the end of epoch 2. The distance between the lap with the global minimum of theta frequency and the end of epoch 2 was not significantly shorter than chance (12.7 laps, p = 0.2246, shuffle test). The gray dashed line represents $95^{th}$ percentile of shuffled data.

e-f. Alternative linear mixed-effect models for the relationship between theta metrics and gain mismatch in landmark-failure sessions. In all models, animal identity was included as a random effect. A fixed effect for gain direction was not included, as all landmark-failure sessions occurred during Gain down conditions. Rat identity is used as a random effect unless specified otherwise. Colors indicate individual animal IDs.

e. (*left*) Full model with random slope and random intercept for dTheta (1 + dTheta | rat). Although the statistical software did not formally flag this model as a singular fit, the model was identified as over-parameterized due to an extreme correlation between random effects (r ≈ -1.0). Consequently, this model was deemed unreliable for interpretation (β = -0.336, CI = [-1.268,0.567], p > 0.05, BIC = 6.0). (*right*) Random-slope-only model for dTheta (0 + dTheta | rat). In contrast to the random-intercept model (main text), which showed a trend, this model shows no significant relationship between dTheta and dH (β = 0.005, CI = [-0.897,0.904], p > 0.05, BIC = 2.4).

f. (*left*) Full model with random slope and random intercept for the theta ratio (1 + theta_ratio | rat). This model showed no significant predictive relationship with dH (β = 0.02, CI = [-0.085,0.130], p > 0.05, BIC = 15.8). (*right*) Random-intercept-only model for the theta ratio (1 | rat). This model resulted in a singular fit, indicated by a zero variance among the random effects, and was therefore considered unreliable for interpretation (β = 0.002, CI = [-0.012,0.017], p > 0.05, BIC = 12.6).

Supplementary Table 1. Parameters values in the Continuous Attractor Neural Network Model

| Parameters | Values |
|---|---|
| Number of neurons: N | 512 |
| Recurrent connection range (Gaussian width): a | 0.4 |
| Input range (Gaussian width): σ | 0.4 |
| Global inhibition strength: k | 5 |
| Adaptation strength: m | 1.1 |
| Allothetic input strength (amplitude): $\alpha_{allo}$ | 0.10 |
| Idiothetic input strength (amplitude): $\alpha_{idio}$ | 0.01 |
| Path-integration alignment distance [m]: D | 2.3253 |
| Path-integration gain: P | Range: [0.3, 1.7] |
| Landmark gain: G | Range: [0.3, 1.7] |
| Gain factor: g | 5 |
| Recurrent connection strength: $J_0$ | 1/5 |
| Animal's speed in landmark frame [m/s]: $V_{allo}$ | 1.45 |
| Time constant of pre-synaptic input (U) [ms]: τ | 3 |
| Time constant of firing rate adaptation (V) [ms]: $\tau_v$ | 144 |
